# A meta-learning approach for genomic survival analysis

**DOI:** 10.1101/2020.04.21.053918

**Authors:** Yeping Lina Qiu, Hong Zheng, Arnout Devos, Olivier Gevaert

## Abstract

RNA sequencing has emerged as a promising approach in cancer prognosis as sequencing data becomes more easily and affordably accessible. However, it remains challenging to build good predictive models especially when the sample size is limited and the number of features is high, which is a common situation in biomedical settings. To address these limitations, we propose a meta-learning framework based on neural networks for survival analysis and evaluate it in a genomic cancer research setting. We demonstrate that, compared to regular transfer-learning, meta-learning is a significantly more effective paradigm to leverage high-dimensional data that is relevant but not directly related to the problem of interest. Specifically, meta-learning explicitly constructs a model, from abundant data of relevant tasks, to learn a new task with few samples effectively. For the application of predicting cancer survival outcome, we also show that the meta-learning framework with a few samples is able to achieve competitive performance with learning from scratch with a significantly larger number of samples. Finally, we demonstrate that the meta-learning model implicitly prioritizes genes based on their contribution to survival prediction and allows us to identify important pathways in cancer.

## 1 Introduction

Cancer is a leading cause of death in the world. Accurate prediction of its survival outcome has been an interesting and challenging problem in cancer research over the past decades. Quantitative methods have been developed to model the relationship between multiple explanatory variables and survival outcome, including fully parametric models (Hosmer et al., 2008; Klein and Moeschberger, 2006) and semi-parametric models such as the Cox proportional hazards model (Cox, 1972). The Cox model makes a parametric assumption about how the predictors affect the hazard function, but makes no assumption about the baseline hazard function itself (Harrell et al., 1982). In most real world scenarios, the form of the true hazard function is either unknown or too complex to model, making the Cox model the most popular method in survival analysis (Kleinbaum and Klein, 2012).

In clinical practice, historically, survival analysis has relied on low-dimensional patient characteristics such as age, sex, and other clinical features in combination with histopathological evaluations such as stage and grade (Louis et al., 2016). With advances in high-throughput sequencing technology, a greater amount of high-dimensional genomic data is now available and more molecular biomarkers can be discovered to determine survival and improve treatment. With the cost of RNA sequencing coming down significantly, from an average of $100M per genome in 2001 to $1k per genome in 2015 (Park and Kim, 2016), it is becoming more feasible to use this technology to prognosticate. Such genomic data often has tens of thousands of variables which requires the development of new algorithms that work well with data of high dimensionality.

To address these challenges, several implementations of regularized Cox models have been proposed (Goeman, 2010; Park and Hastie, 2007; Wong et al., 2018). A regularized model adds a model complexity penalty to the Cox partial likelihood to reduce the chance of overfitting. More recently, the increasing modeling power of deep learning networks has aided in developing suitable survival analysis platforms for high dimensional feature spaces. For example, autoencoder architectures have been employed to extract features from genomic data for liver cancer prognosis prediction (Chaudhary et al., 2018). The Cox model has also been integrated in a neural network setting to allow greater modeling flexibility (Cheerla and Gevaert (2019); Ching et al. (2018); Luck et al. (2017); Yousefi et al. (2017).

In studying a specific rare cancer’s survival outcome, one interesting problem is whether it is possible to make use of the abundant data that is available for more common relevant cancers and leverage that information to improve the survival prediction. This problem is commonly approached with transfer-learning (Pratt, 1993), where a model which has been trained on a single task (e.g., 1 or more abundant cancers) is used to fine-tune on a related target task (rare cancer). In survival analysis, transfer-learning has shown to significantly improve prediction performance (Li et al., 2016). Deep neural networks used to analyze biomedical imaging data can also take advantage of information transfer from data in other settings. For example, multiple studies show that convolutional neural networks pretrained on ImageNet data can be used to build performant survival models with histology images (Deng et al., 2009; Mobadersany et al., 2018).

In this context, meta-learning is an area in deep learning research that has gained much attention in recent years which addresses the problem of “learning to learn” (Finn et al., 2017; Vilalta and Drissi, 2002). A meta-learning model explicitly learns to adapt to new tasks quickly and efficiently, usually with a limited exposure to the new task environment. Such a framework may potentially adapt better than the traditional transfer-learning setting where there is no explicit adaptation incorporated in the pre-training algorithm. This problem setting with limited exposure to a new task is also known as few-shot learning: learn to generalize well, given very few examples (called *shots*) of a new task (Li et al., 2016). Recent advances in meta-learning have shown that, compared to transfer-learning, it is a more effective approach to few-shot classification (Chen et al., 2017; Devos and Grossglauser, 2019), regression (Finn et al., 2017), and reinforcement learning (Duan et al., 2016; Finn et al., 2017). In this study, we propose a meta-learning framework based on neural networks for survival analysis applied in a cancer research setting. Specifically, for the application of predicting survival outcome, we demonstrate that our method is a preferable choice compared to regular transfer-learning pre-training and other competing methods on three cancer datasets when the number of training samples from the specific target cancer is very limited. Finally, we demonstrate that the meta-learning model implicitly prioritizes genes based on their contribution to survival prediction and allows us to uncover biological pathways associated with cancer survival outcome.

## 2 Materials and methods

### 2.1 Datasets

We use the RNA sequencing data from The Cancer Genome Atlas (TCGA) pan-cancer datasets (Tomczak et al., 2015). We remove the genes with NA values and normalized the data by log transformation and z-score transformation. The feature dimension is 17176 genes after preprocessing. The data contains 9707 samples from 33 cancer types. The outcome is the length of survival time in months. 78% of the patients are censored, which means that the subject leaves the study before an event occurs or the study terminates before an event occurs to the subject.

### 2.2 Meta-learning

Our proposed survival prediction framework is based on a neural network extension of the Cox regression model that relies on semi-parametric modeling by using a Cox loss (Ching et al., 2018). The model consists of two modules: the feature extraction network and the Cox loss module (Figure 1). We use a neural network with two hidden layers to extract features from the RNA sequencing data input, which yields a lower dimensional feature vector for each patient. The features are then fed to the Cox loss module, which performs survival prediction by doing a Cox regression with the features as linear predictors of the hazard (Cox, 1972). The parameters of the Cox loss module *β* are optimized by minimizing the negative of the partial log-likelihood:

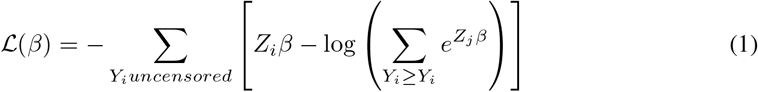

where *Y_i_* is the survival length for patient *i, Z_i_* is the extracted feature for patient *i*, and *β* is the coefficient weight vector between the feature and the output. Since *Z_i_* is the output of the feature extraction module, it can be further represented by:

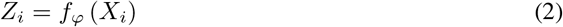

where *X_i_* is the input predictor of patient i, *f* denotes a nonlinear mapping the neural network learns to extract features form the predictor, and *φ* denotes the model parameters including the weights and biases of each neural network layer. The feature extraction module parameters *φ* and Cox loss module parameters *β* are jointly trained in the model. For convenience in the following discussion we denote the combined parameters as *θ*.

**Figure 1:**
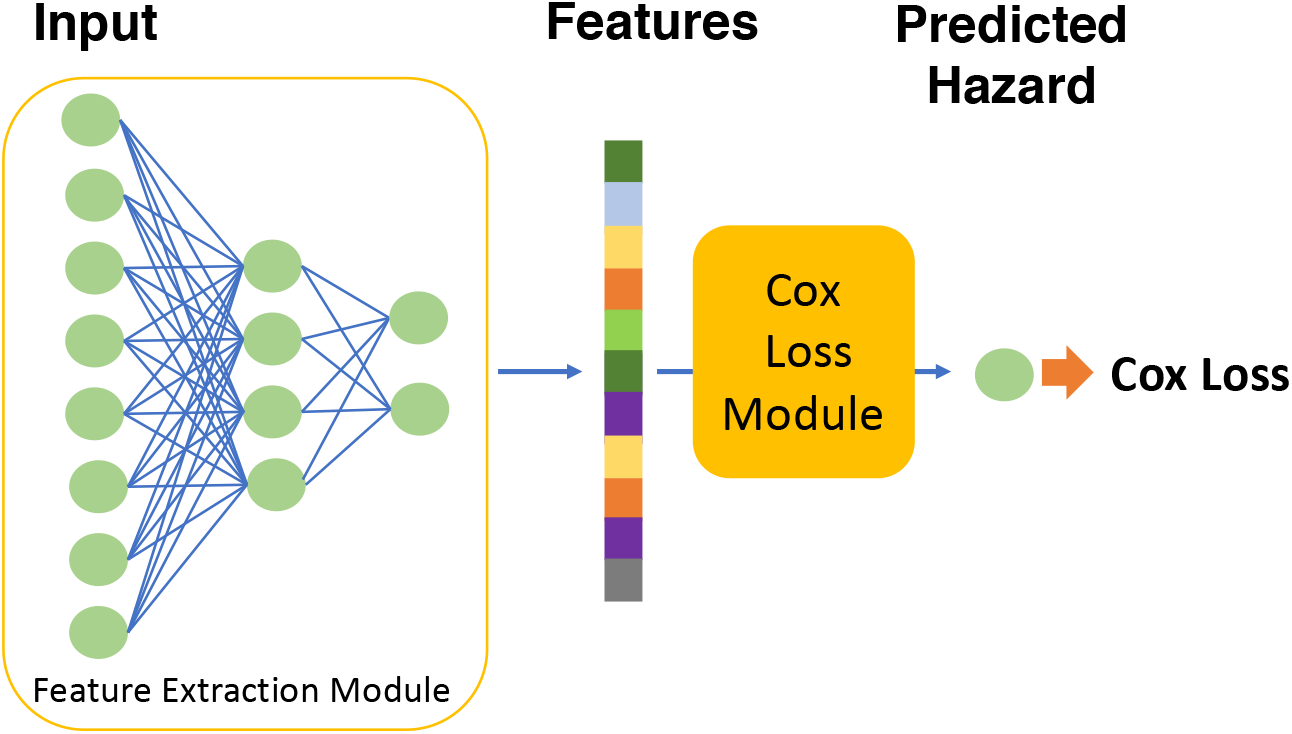
Schematic showing the survival prediction model architecture

The optimization of parameters *θ* consists of two stages: a meta-learning stage, and a final learning stage. The meta-learning stage is the key process, where the model aims to learn a suitable parameter initialization for the final learning stage, so that during final learning the model can adapt efficiently to previously unseen tasks with a few training samples (Nichol et al., 2018). In order to reach the desired intermediate state, a first order gradient-based meta-learning algorithm is used to train the network during the meta-learning stage (Finn et al., 2017; Nichol et al., 2018).

Specifically, at the beginning of meta-learning training, the model is randomly initialized with parameter *θ*. Consider that the training samples for the meta-learning stage consist of n tasks *T_τ_*, *τ* = 1,2 … *n*. A task is defined as a common learning theme shared by a subgroup of samples. Concretely, these samples come from a distribution on which we want to carry out a classification task, regression task or reinforcement learning task. The algorithm continues by sampling a task *T_τ_* and using samples of *T_τ_* to update the inner-learner. The inner-learner learns *T_τ_* by taking k steps of stochastic gradient descent (SGD) and updating the parameters to 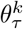:

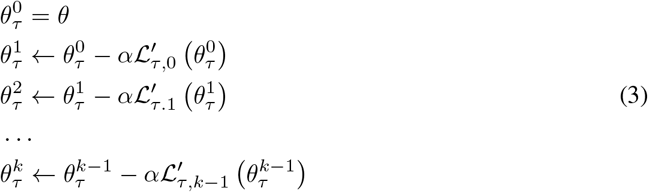

where 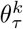 is the model parameter at step k for learning task *τ*, 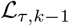 is the loss computed on the k^th^ minibatch sampled from task *τ*, 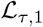 is the loss computed on the second minibatch sampled from task *τ* and so on. The ‘prime’ symbol denotes differentiation, and *α* is the inner learner step size. Note that this learning process is the separate for all tasks, starting from the same initialization *θ*.

After an arbitrary *m* number of tasks are independently learnt by the above k-step SGD process, and obtaining 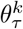, *τ* = 1, 2,… m, we make one update across all tasks with the meta-learner to get a better initialization *θ*:

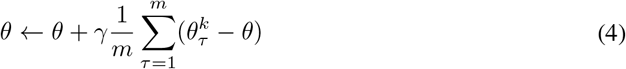

where *γ* is the learning step of the meta-learner. The term 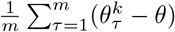 can be considered as a gradient, so that for example a popular optimization algorithm like Adam (Kingma and Ba, 2014) can be used by the meta-learner to self-adjust learning rates for each parameter. The entire process of inner-learner update and meta-learner update is repeated until a chosen maximum number of meta-learning epochs is reached. This algorithm is shown to encourage the gradients of different minibatches of a given task to align in the same direction, thereby improving generalization and efficient learning later on (Nichol et al., 2018).

In the final learning stage, the model is provided with a few-sample dataset of a new task. First, the model is initialized with the meta-learnt parameters *θ*, which are then fine-tuned with the new task training data to 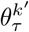 and finally the fine-tuned model is evaluated with testing data from the new task. The training procedure of final learning does not require a special algorithm, and can be conducted with regular mini-batch stochastic gradient descent. This final learning stage is equal to the inner-learning loop for a single task in Equation (3) without any outer loop.

### 2.3 Experimental setup

In order to assess the meta-learning method’s performance, we compare it with several alternative training schemes based on the same neural network architecture: regular *pre-training, combined learning*, and *direct learning*. First, meta-learning initially learns general knowledge from a dataset containing tasks that are relevant but not directly related to the target, and then learns task-specific knowledge from a very small target task dataset. We define the first dataset as the “multi-task training data”, and the second as the “target task training data”. Secondly, regular pre-training also has a two-stage learning process on the same datasets, but unlike meta-learning without explicitly focusing on learning to reach an initialization that is easy to adapt to new tasks. Thirdly, combined learning does not involve a two-stage learning process, but also leverages knowledge from the relevant tasks by combining the multi-task training data and the target task training data together in one dataset to train a prediction model. Direct learning on the other hand, only uses the target task training samples. To illustrate the effectiveness of the methods with few samples, we consider three cases of direct learning: a large sample size, a medium sample size, and a small sample size which is the same size as the “target task training data” used for the other methods (i.e. regular pre-training, combined learning and meta-learning) (Figure 2).

**Figure 2:**
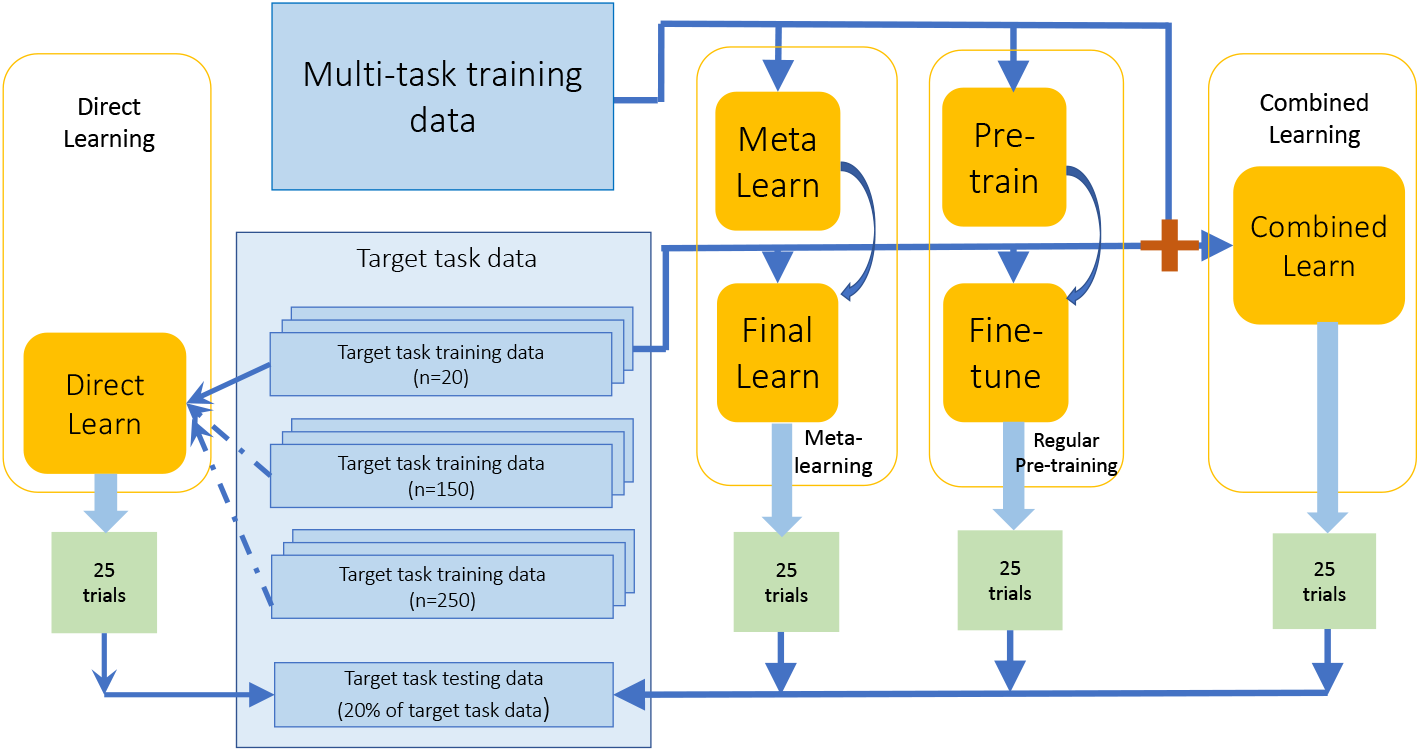
Data flow for meta-learning, regular pre-training, and combined learning frameworks

In our experiments, the “multi-task training data” is the pan-cancer RNA sequencing data containing samples from any cancer sites except one cancer site that we define as the target cancer site. The associated target cancer data is considered as the “target task data”. This target task dataset is split into training data (80%) and testing data (20%), stratified by disease sub-type and censoring status.

For meta-learning, regular pre-training, and combined learning we will not use all 80% of the training set for the target task, as we want to assess the performances when the algorithm is exposed to only a small number of target task training samples. Therefore, we will randomly draw 20 samples from the training dataset as one “target task training data”. We choose a small sample size of 20 because it is a possible case in real life situations where the target task is the study of rare diseases (Hee et al., 2017), or where new technologies are used to produce data which only have the capacity to produce a small sample. For direct learning, we randomly draw three different sizes from the training datasets to form training data, 20 for the small size, 150 for the medium size, and 250 for the large size. All methods are evaluated on the common testing data of the target task.

Finally, as a linear baseline, we use the combined learning training sample (multi-task training data and target task training data) to train a linear cox regression model. We conduct 25 experiment trials for each method, where each trial is trained with a randomly drawn “target task training dataset” as described above.

### 2.4 Evaluation

We evaluate the survival prediction model performances with two commonly used evaluation metrics: the concordance index (C-index) (Harrell et al., 1982) and the integrated Brier score (IBS) (Brier, 1950). Fistly, the C-index is a standard performance measure of a model’s predictive ability in survival analysis. It is calculated by dividing the number of all pairs of subjects whose predicted survival times are correctly ordered, by the number of pairs of subjects that can possibly be ordered. A pair cannot be ordered if the earlier time in the pair is censored or both events in the pair are censored. A C-index value of 1.0 indicates perfect prediction where all the predicted pairs are correctly ordered, and a value of 0.5 indicates random prediction. Secondly, the integrated Brier score is used to evaluate the error of survival prediction and is represented by the mean squared differences between observed survival status and the predicted survival probability at a given time point. The IBS provides an overall calculation of the model performance at all available times. An IBS value of 0 indicates perfect prediction, while 1 indicates entirely inaccurate prediction.

We select cancer sites from TCGA with the following two inclusion criteria: (1) a minimum of 450 samples, providing enough training samples for different benchmarking training schemes and (2) a minimum of 30% non-censoring samples, enabling more accurate evaluation than more heavily censored cohorts. This results in the following cancers: glioma, including glioblastoma (GBM) and low-grade-glioma (LGG); non-small cell lung cancer, including lung adenocarcinoma (LUAD) and lung squamous cell carcinoma (LUSC), and head-neck squamous cell carcinoma (HNSC). In addition, these three types of cancers are of clinical interest, because gliomas are the most common type of malignant brain tumor, and lung cancer is the deadliest cancer in the world (Ceccarelli et al., 2016; Herbst and Lippman, 2007). HNSC, on the other hand, is a less widely studied type of cancer, which nonetheless attracted growing attention in the recent decade since the release of the publicly available largest dataset in HNSC by TCGA (Brennan et al., 2017; Tonella et al., 2017).

We evaluate the C-index and IBS in 25 experimental trials for each method.

### 2.5 Hyper-parameter selection

To avoid overfitting, we do not conduct a separate hyper-parameter search for each of the cancer datasets. Instead, we search for hyper-parameters on one type of cancer and apply the chosen parameters to all experiments. We select the largest cancer cohort, glioma, and use 5-fold crossvalidation for hyper-parameter selection. For each given set of hyper-parameters, we average the results from five validation sets (each is 20% of training data). Since there is similarity in the algorithm between methods (combined learning, direct learning, and regular pre-training), we share hyper-parameters between experiments when it makes sense, as detailed below.

All methods use the same neural network architecture with two hidden fully connected layers of size 6000 and 2000, and an output fully connected feature layer of size 200. Each layer uses the ReLU activation function (Nair and Hinton, 2010). Initially we experiment with 4 different structures: 1 or 2 hidden layers with feature size of 200 or 50, respectively. We chose the optimal structure detailed before and use it as the architecture for all methods in our subsequent discussion.

For the regular pre-training model, we search for hyper-parameters for the pre-training stage and fine-tuning stage separately. For both stages, we test the mini-batch gradient descent and Adam optimizers, and determine learning rates with grid search on a grid of [.1, .05, .01, .005, .001] for SGD and a grid of [.001, .0005, .0001,.00005, .00001] for Adam. We test batch sizes of 50, 100, 200, and 800 for pre-training. The selected parameters for the pre-train stage are: an SGD optimizer with learning rate of .001, L2 regularization scale of 0.1 and batch size of 800. The selected parameters for the fine-tune stage are: an SGD optimizer with learning rate of .001, L2 regularization scale of 0.1, and batch size of 20 which is the size of each target cancer training dataset. For the combined learning model and direct learning model, since the algorithm is very similar to the regular pre-training model’s pre-train stage, we use the same parameters selected for the pre-train. The batch sizes for direct learning is half of the size of training data.

For the meta-learning model, we search for hyper-parameters for the meta-learning only. For the final learning stage, we use the same hyper-parameters as in the fine-tune stage of the regular pre-training model, as both methods can use similar algorithms in the last stage of training. From our previous discussion, in the meta-learning stage an SGD optimizer and an Adam optimizer are suitable for the inner learner and meta-learner respectively. For the learning rates, we perform grid search on a grid of [.1, .05, .01, .005, .001] for SGD and a grid of [.001, .0005, .0001,.00005, .00001] for Adam. Batch size is searched from [50, 100, 200, 800], the number of tasks for averaging one meta-learner update is searched from [5, 10, 20], and the number of gradient descent steps for the inner-learner is searched from [3, 5, 10, 20]. The selected parameters for the meta-learning stage are shown in Table 1.

**Table 1:** Selected hyper-parameters for meta-learning’s meta-learning stage

| Hyper-parameter | Value |
| --- | --- |
| Task-level optimizer | SGD |
| Task-level learning rate | 0.01 |
| Task-level gradient steps | 5 |
| Task-level Batch size | 100 |
| Meta-level optimizer | ADAM |
| Meta-level learning rate | 0.0001 |
| Meta-level tasks batch size | 10 |
| L2 regularization scale | 0.1 |

Finally, in order to evaluate the effect of fluctuations of the meta-learning hyper-parameters, and ensure that our results reflect the average performance over fluctuations, we conduct a series of tests on the validation data where in each experiment we vary one of the five unique meta-learning hyper-parameters from the chosen value by tuning it up or down by one grid, obtaining ten sets of varied hyper-parameters. We do 5-fold cross-validation for each set of varied hyper-parameters and compute the concordance index from the resulting fifty experiments. We also do fifty random experiments using the selected hyper-parameters and compare the average results of varied versus selected hyper-parameters. We conduct a two-sample t-test on the two results, and conclude that the results obtained by varied parameters do not have a significant difference from the results obtained by the chosen parameters (mean concordance index difference of 0.005 with p value 0.50). Therefore, our results are robust with respect to fluctuations of the hyper-parameters and our conclusions are not based on excessive hyper-parameter tuning.

### 2.6 Interpretation of the genes prioritized by the meta-learning model

We apply *risk score backpropagation* (Yousefi et al., 2017) to the meta-learned models to investigate the feature importance of genes for each of the three target cancer sites. For a given sample, each input feature is assigned a risk score by taking the partial derivatives of the risk with respect to the feature. A positive risk score with high absolute value means the feature is important in poor prediction (high risk), and a negative risk score with high absolute value means the feature is important in good prediction (low risk). The features are ranked by the average of risk score across all samples.

Two approaches were adopted for annotating the genes with ranked risk scores generated by the meta-learning model. Firstly, the top 10% high-risk genes (genes with positive risk scores) and the top 10% low-risk genes (genes with negative risk scores) from each cancer type were subjected to gene set over-representation analysis, by comparing the genes against the gene sets annotated with well-defined biological functions and processes. We model the association between the genes and each gene set using a hypergeometric distribution and Fisher’s exact test. Secondly, instead of arbitrary thresholding in the first approach, all the genes, together with their ranked risk scores were incorporated in the gene set enrichment analysis with the fgsea R package (Sergushichev, 2016) which calculates a cumulative gene set enrichment statistic value for each gene set. The gene set databases used in this analysis include Kyoto Encyclopedia of Genes and Genomes (KEGG) (Kanehisa and Goto, 2000), The Reactome Pathway Knowledgebase (Croft et al., 2014) and WikiPathways (Slenter et al., 2018).

## 3 Results

### 3.1 Meta-learning outperforms regular pre-training and combined learning

For all of the target cancer sites, meta-learning achieves similar or better performance than regular pre-training or combined learning (Figure 3; Table 2). For the glioma cohort, the mean C-index for meta-learning is 0.86 (0.85-0.86 95% CI), compared to 0.84 (0.83-0.85 95% CI) for regular pre-training and 0.81 (0.81-0.82 95% CI) for combined training. For the lung cancer cohort, the mean C-index is 0.65 (0.65-0.66 95% CI) for meta-learning, 0.60 (0.58-0.61 95% CI) for regular pre-training, and 0.62 (0.62-0.63 95% CI) for combined training. For the HNSC cohort, the result is 0.61 (0.59-0.63 95% CI) for meta-learning, 0.59 (0.57-0.61 95% CI) for regular pre-training and 0.62 (0.61-0.64 95% CI) for combined training. Note that, the variance of the meta-learning results across 25 random trials also tends to be the smallest, which is most observable for the lung cancer and glioma cohorts. In addition, each of these multi-layer neural networks also shows better performance on average than a linear baseline model. The linear baseline model achieves a C-index of 0.61 for lung cancer (0.60-0.62 95% CI), 0.77 for glioma (0.74-0.80 95% CI), and 0.59 for HNSC (0.58-0.62 95% CI).

**Figure 3:**
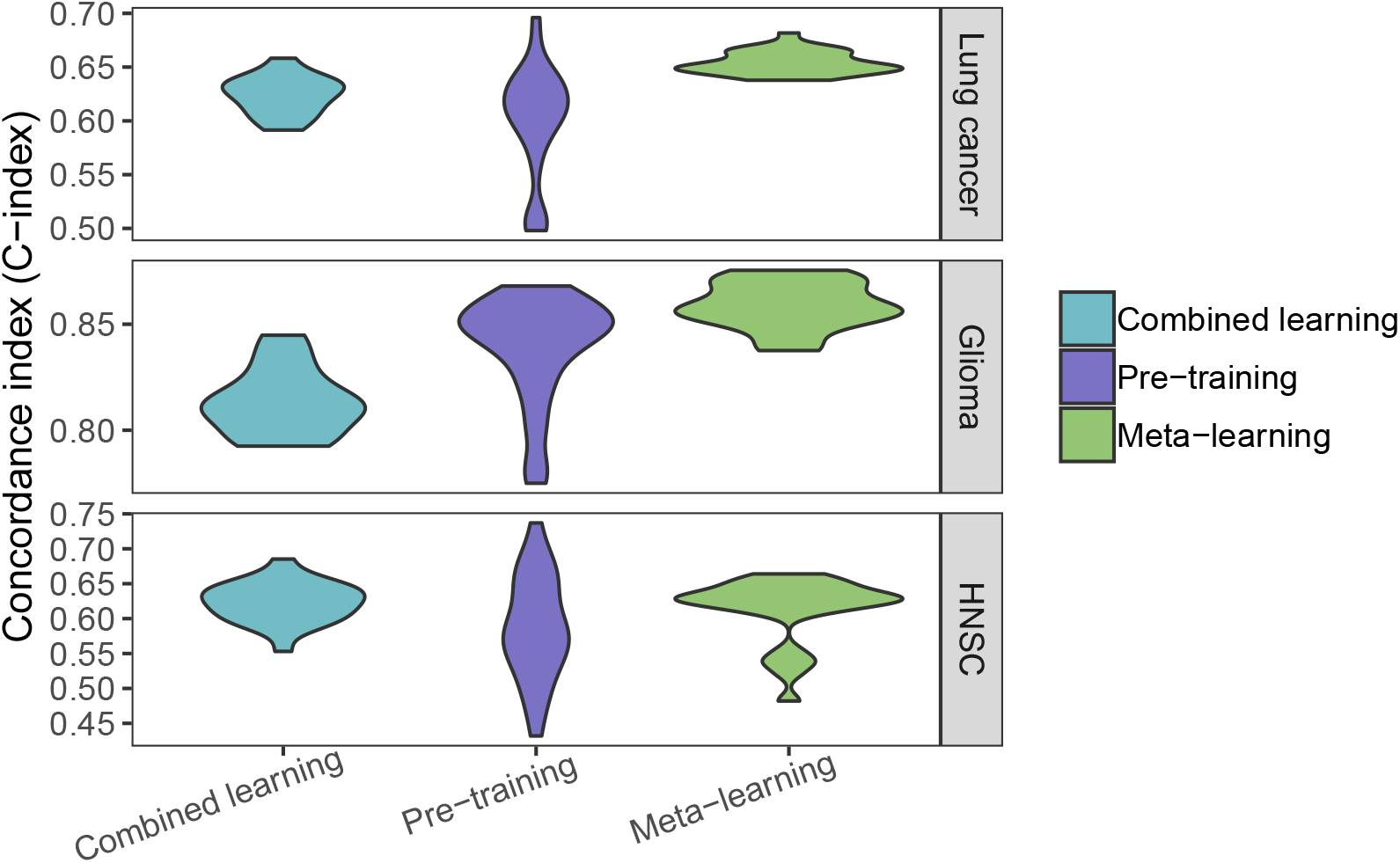
C-Index for target cancer survival prediction, comparing combined learning, regular pre-training and meta-learning

**Table 2:**
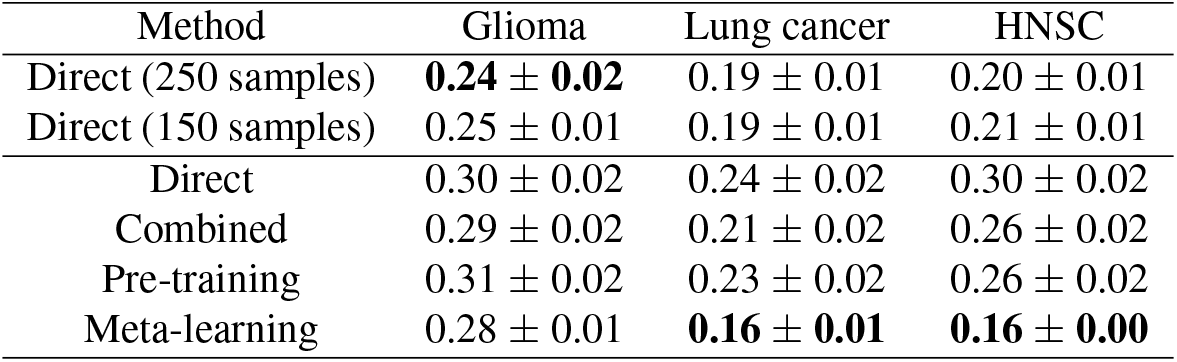
Integrated Brier scores (IBS) with 95% confidence intervals (n=X trials) for target cancer survival prediction with 20 samples, unless specified otherwise. Lower value is better. Best performing method in bold.

| Method | Glioma | Lung cancer | HNSC |
| --- | --- | --- | --- |
| Direct (250 samples) | **0.24 ± 0.02** | 0.19 ± 0.01 | 0.20 ± 0.01 |
| Direct (150 samples) | 0.25 ± 0.01 | 0.19 ± 0.01 | 0.21 ± 0.01 |
| Direct | 0.30 ± 0.02 | 0.24 ± 0.02 | 0.30 ± 0.02 |
| Combined | 0.29 ± 0.02 | 0.21 ± 0.02 | 0.26 ± 0.02 |
| Pre-training | 0.31 ± 0.02 | 0.23 ± 0.02 | 0.26 ± 0.02 |
| Meta-learning | 0.28 ± 0.01 | **0.16 ± 0.01** | **0.16 ± 0.00** |

### 3.2 Meta-learning achieves competitive predictive performance compared to direct learning

Next, we compare our meta-learning approach with regular direct learning on the target task training samples with different cohort sizes. The performance of direct learning drops significantly when the number of training samples decreases from 250 to 20, which is anticipated because a great amount of information is lost and the model can hardly learn well. However, meta-learning and pretraining can compensate for such lack of information by transferring knowledge from the pan-cancer data explicitly and implicitly, respectively. We show that meta-learning achieves similar or better prediction performance than large-sample direct training in lung cancer and HNSC, and reaches comparable performance with medium-sample direct training in glioma (Figure 4; Table 2). For the lung cancer cohort, the mean C-index is 0.57 (0.56-0.58 95% CI) for large sample direct learning, 0.54 (0.52-0.56 95% CI) for medium sample direct learning, 0.53 (0.50-0.55 95% CI) for small sample direct learning, and 0.65 (0.65-0.66 95% CI) for meta-learning. For the glioma cohort, the mean C-indices for large sample, medium sample and small sample direct learning are 0.86 (0.86-0.87 95% CI), 0.85 (0.85-0.86 95% CI), and 0.82 (0.81-0.84 95% CI) respectively, and for meta-learning the mean C-index is 0.86 (0.85-0.86 95% CI). For the HNSC cohort, the mean C-index is 0.62 (0.60-0.64 95% CI) for large sample direct learning, 0.57 (0.54-0.59 95% CI) for medium sample direct learning, 0.53 (0.49-0.56 95% CI) for small sample direct learning, and 0.61 (0.59-0.63 95% CI) for meta-learning. Thus, in all three cancer sites, meta-learning reaches competitive performances as large sample direct learning and can outperform it in certain cases.

**Figure 4:**
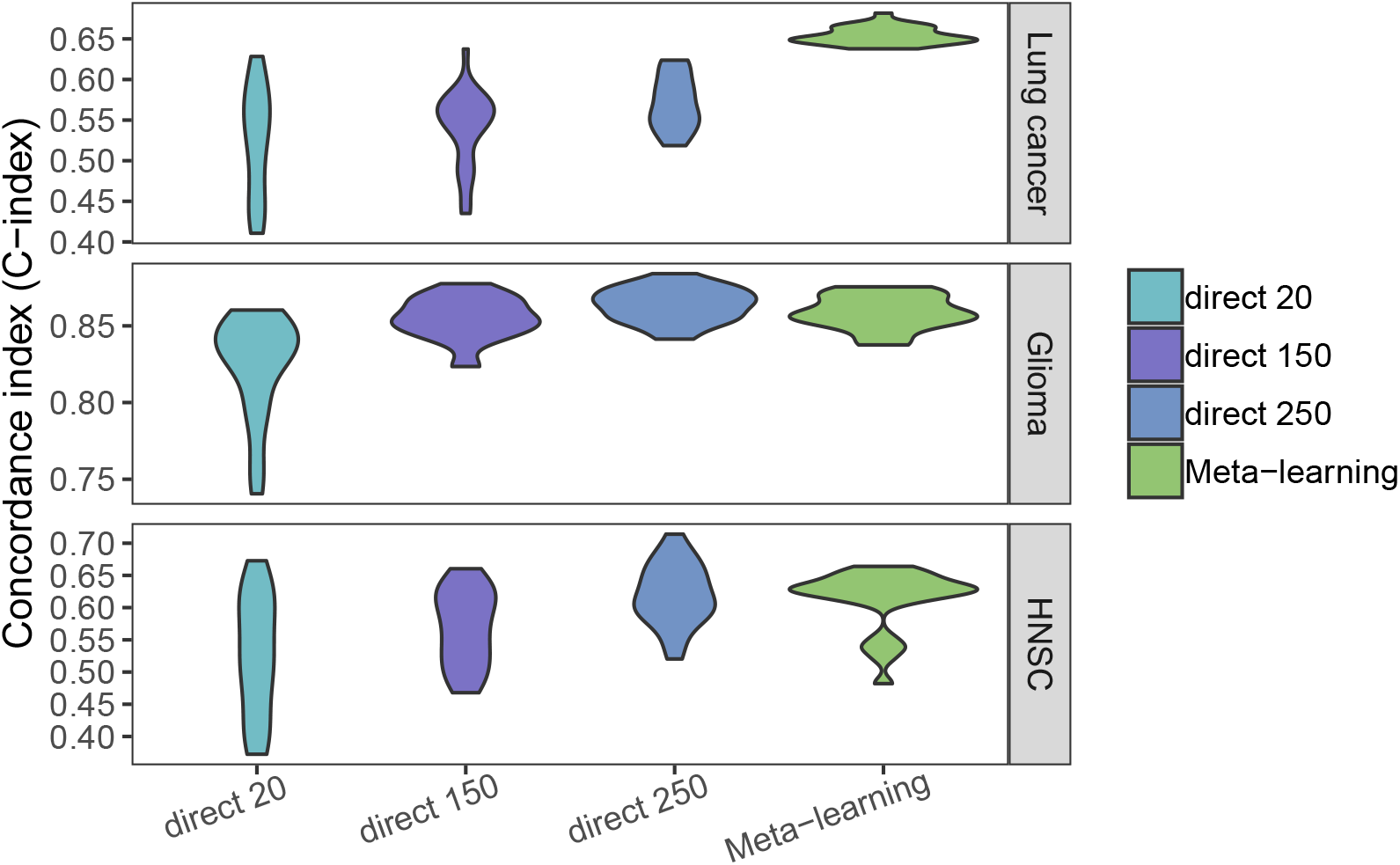
C-Index for target cancer survival prediction, comparing direct learning with large (250), medium (150) and small (20) size samples and meta-learning.

### 3.3 Risk score ranked genes are enriched in key cancer pathways

Next, we investigated for each cancer site what genes are most important in the meta-learning model (Figure 5, Supplementary Tables S1-S6). In gliomas, the high-risk genes are associated with viral carcinogenesis (p value 0.002), Herpes simplex infection (p value 0.007), cell cycle (p value 0.03), apoptosis (p value 0.03), DNA damage response (p value 0.04), all of which are also enriched in gene set enrichment analysis with positive enrichment scores (Figure 5a, Table S2). The low-risk genes are associated with HSF1 activation (p value 0.02) which is involved in hypoxia pathway, and tryptophan metabolism (p value 0.04), the latter is also enriched in gene set enrichment analysis with negative enrichment score. Tryptophan catabolism has been increasingly recognized as an important microenvironmental factor in anti-tumor immune responses (Platten et al., 2012) and it is a common therapeutic target in cancer and neurodegeneration diseases (Platten et al., 2019).

**Figure 5:**
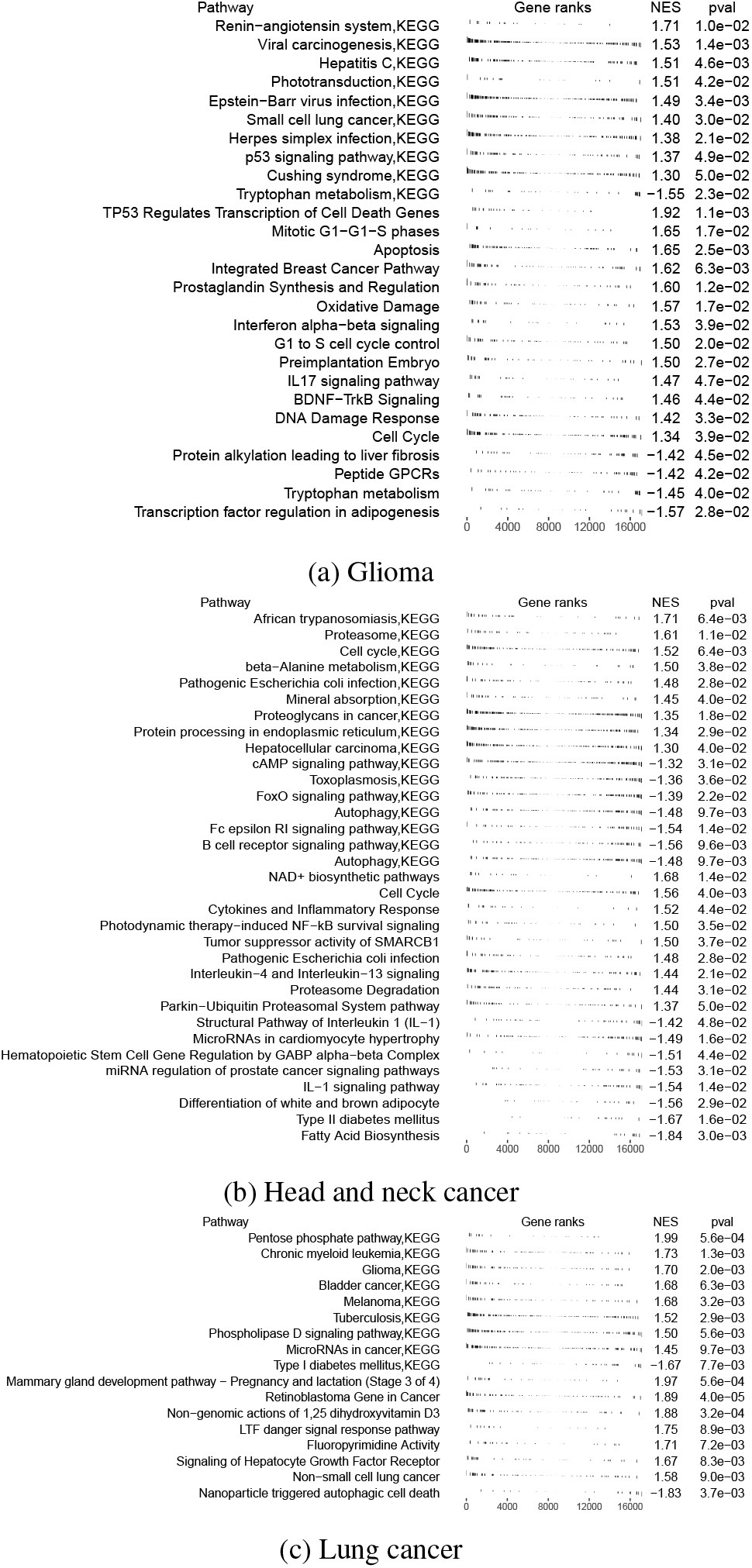
Gene set enrichment analysis of the ranked gene list in (a) Glioma (b) Head and neck cancer (c) Lung cancer. The gene set databases used in this analysis included Kyoto Encyclopedia of Genes and Genomes (KEGG) and WikiPathways. Pval, enrichment p-value; NES, normalized enrichment score. In (a) and (b) the enrich pathways with p value below 0.05 were displayed. In (c) the enrich pathways with p value below 0.01 were displayed. Detailed information can be found in supplementary tables S2, S4 and S6.

In head and neck cancer, the high-risk genes are associated with PTK6 signaling (p value 0.01), which regulates cell cycle and growth, and cytokines and inflammatory response (p value 0.009). The low-risk genes are associated with autophagy (p value 0.02), which is also enriched in gene set enrichment analysis. Other enriched pathways include B cell receptor signaling pathway, cell cycle, and interleukin 1 signaling pathway (Figure 5b, Table S4). Interleukin 1 is an inflammatory cytokine which plays a key role in carcinogenesis and tumor progression (Mantovani et al., 2018).

In lung cancer, the top high-risk genes are associated with "non-small cell lung cancer" pathway (p value 0.01), tuberculosis (p value 0.008), Hepatitis B and C virus infection (p value 0.03), and many pathways implicated previously in cancer. These pathways are also enriched in gene set enrichment analysis (Figure 5c, Table S6). Pulmonary tuberculosis has been shown to increase the risk of lung cancer (Wu et al., 2011; Yu et al., 2011). The low-risk genes are associated with energy metabolism (p value 0.03), ferroptosis (p value 0.037) and AMPK signaling pathway (p value 0.046), all related to energy metabolism, particularly lipid metabolism. AMPK signaling pathway activation by an AMPK agonist was shown to suppresses non-small cell lung cancer through inhibition of lipid metabolism (Chen et al., 2017). AMPK signaling and energy metabolism are also enriched in gene set enrichment analysis (Figure 5C). Other enriched pathways include Notch signaling, interleukin signaling, ErbB signaling, and signaling pathways regulating pluripotency of stem cells.

## 4 Discussion

Previous studies have shown that when analyzing high-dimensional genomic data, deep learning survival models can achieve comparable or superior performance compared to other methods (e.g., Cox elastic net regression, random survival forests) (Kim et al., 2019). However, the performance of deep learning is often limited by the relatively small amount of available data (Yousefi et al., 2017). To address this issue, our work investigates different deep learning paradigms to improve the performance of deep survival models, especially in the setting of small size training data.

In previous studies the most common way to build deep survival models is to train neural networks with a large number of target task training samples from scratch, a process we call direct training. Direct training with a large sample size (e.g. *n* = 250) can thus be considered as a baseline. As expected, the performance of direct training drops when the number of training samples decreases (e.g. from *n* = 150 to *n* = 20). On the other hand, combined learning, regular pre-training, and meta-learning all leverage additional data from other sources, thereby enabling them to achieve better performances when the training sample size is small. We use a small number of target cancer site training samples (e.g. *n* = 20) with these methods and investigate their performance.

It is important to note that combined learning, regular pre-training and meta-learning are exposed to exactly the same information, but differ only in their algorithms. Combined learning is a one-stage learning process, whereas pre-training and meta-learning are two-stage learning methods. Metalearning shows better predictive performance than combined or regular pre-training, indicating that it is able to adapt to a new task more effectively due to the improved optimization algorithm targeting the few-sample training environment.

It has been shown that methods which use only target task data (direct learning with different size samples) and methods which use additional information (combined, pre-training and meta-learning) perform differently, and one type of approach may be better than the other on different cancer sites. For example, on glioma, direct learning tends to do better overall; whereas on lung cancer, the other methods outperform direct learning. This may be due to that fact that the amount of information that can be learnt from related data versus from the target data is different for each cancer site. If there is significant information within the target cancer samples alone, then direct training will be more effective than learning from other cancer samples. On all three cancer sites we observe that meta-learning achieves similar or better performance than medium-size direct training, and outperforms large-size direct training in some cases. However, the advantage of meta-learning may not generalize to every cancer site. Certain cancers may have very unique characteristics so that transfer of information from other cancers may not help in prediction regardless of improved adaptivity. For the three cancer sites, the affinity of each target cancer to other types of cancers in the pan-cancer data aids the performance of meta-learning, which efficiently transfers the information from other cancers to the target cancers. On the other hand, some cancers are more dissimilar from other cancer sites which makes information transfer difficult. For example, for another cancer site, kidney cancer, specifically kidney renal clear cell carcinoma (KIRC) and kidney renal papillary cell carcinoma (KIRP), both meta-learning and pre-training do not produce good survival prediction. This can be visualized in Figure S1, comparing the affinity between different target cancers with the rest of the cancers on a t-distributed stochastic neighbor embedding (t-SNE) graph. Therefore, in order for meta-learning to achieve good performance, the related tasks training data need to contain a reasonable amount of transferable information to the target task.

The performance of meta-learning can be explained by the learned learning algorithm at the metalearning stage where the model learns from related tasks. We further investigate how to optimize meta-learning performance. We examine results from two sampling approaches when forming one task, where we either draw samples only from one cancer, or draw samples from multiple types of cancer. It is a more natural choice to consider each cancer type as a separate task, but we found that the latter leads to improved performance. To explain this improvement, we examine the gradient of the meta-learning loss function. It can be shown that the gradient of the loss function contains a term that encourages the gradients from different minibatches for a given task to align in the same direction (Appendix A). If the two minibatches contain samples from the same type of cancer, their gradient might already be very similar and thus this higher order term would not have a large effect. On the other hand, if the second minibatch contains samples from a different type of cancer than the first, the algorithm will learn something that is common to both of them and thereby help to improve generalization.

The gene set enrichment analysis results validate our model for prioritizing the genes for survival predication. In the three cancer types investigated, the resulting gene lists are enriched in key pathways in cancer including cell cycle regulation, DNA damage response, cell death, interleukin signaling, and NOTCH signaling pathway, etc.

Apart from the well-recognized cancer pathways, our results also reveal potential players affecting cancer development and prognosis, that are not well-studied yet. Viruses have been linked to the carcinogenesis of several cancers, including human papilloma virus in cervical cancer, hepatitis B and C viruses in liver cancer, and Epstein-Barr virus in several lymphomas and nasopharyngeal carcinoma (Martin and Gutkind, 2008). Our results further suggest that viruses might also play a role in glioma and lung cancer, where the high-risk genes are enriched in several viral carcinogenesis pathways. In gliomas, the enriched pathways that are unfavorable for survival include Epstein-Barr virus and herpes simplex infection. In lung cancer, both hepatitis B and C virus infection pathways are enriched. This suggests that that hepatotropic viruses may affect the respiratory system, including the association with lung cancer. For example, hepatitis B virus infection has been associated with poor prognosis in patients with advanced non-small cell lung cancer (Peng et al., 2015). The role of Epstein-Barr virus in gliomagenesis have also been studied but the results remain inconclusive (Akhtar et al., 2018). Whether these viruses do play a role in carcinogenesis and further affect cancer prognosis, or the association we observed reflects an abnormal immune system that is unfavorable for the survival of cancer patients remains to be investigated.

As for the enriched pathways that are favorable for cancer survival, we identified pathways related to metabolism, in particular, lipid metabolism, in all the three cancer types investigated. In glioma, the top enriched pathway favorable for cancer survival is adipogenesis regulation. In head and neck cancer, differentiation of adopocyte and fatty acid biosynthesis are top enriched favorable pathways. In lung cancer, ferroptosis and AMPK signaling pathway are both related to energy metabolism. Ferroptosis is a process driven by accumulated iron-dependent lipid ROS that leads to cell death. Small molecules-induced ferroptosis has a strong inhibition of tumor growth and enhances the sensitivity of chemotherapeutic drugs, especially in drug resistance (Lu et al., 2018). AMPK plays a central role in the control of cell growth, proliferation and autophagy through the regulation of mTOR activity and lipid metabolism (Chen et al., 2017; Han et al., 2013). The link between cancer and metabolism is worth investigating in future studies.

## 5 Conclusion

In survival analyses one problem that researchers have encountered is the insufficient amount of training samples for machine learning algorithms to achieve good performances. We address this problem by adapting a meta-learning approach which learns effectively with only a small number of target task training samples. We show that the meta-learning framework is able to achieve similar performance as learning from a significantly larger number of samples by using an efficient knowledge transfer. Moreover, in the context of limited training sample exposure, we demonstrate that this framework achieves superior predictive performance over both regular pre-training and combined learning methods on two types of target cancer sites. Finally, we show that meta-learning models are interpretable and can be used to investigate biological phenomena associated with cancer survival outcome.

The problem of small data size may be a limiting factor in many biomedical analyses, especially when studies are conducted with data that is expensive to produce, or in the case of multi-modal data (Cheerla and Gevaert, 2019). Our work shows the promise of meta-learning for biomedical applications to alleviate the problem of limited data. In future work, we intend to extend this approach to analysis with medical imaging data, such as histopathology data and radiology data, for building predictive models on multi-modal data with limited sets of patients.

## 6 Acknowledgements

AD acknowledges funding from the European Union’s Horizon 2020 research and innovation programme under the Marie Skłodowska-Curie grant agreement No. 754354.

## Appendix A

We compute the gradient of the meta-learning loss function. Suppose that, for each task *T_τ_*, the inner-learner takes 2 steps of stochastic gradient descent and updates the parameters to 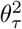. From Equation (3), we can write the gradient using a Taylor expansion:

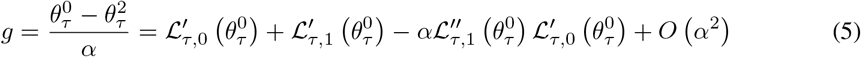

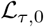 is the loss computed on the first minibatch sampled from task *τ*, and 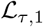 is the loss computed on the second minibatch sampled from task *τ*. The expectation of the first two terms in Equation (5) corresponds to the gradient of expected loss, and the expectation of the third term can be written as:

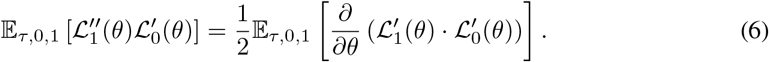

This term increases the inner product of the gradient of the first minibatch and the gradient of the second minibatch, which means it encourages the gradients from different minibatches for a given task to align in the same direction.

## Supplementary information

**Figure S1:**
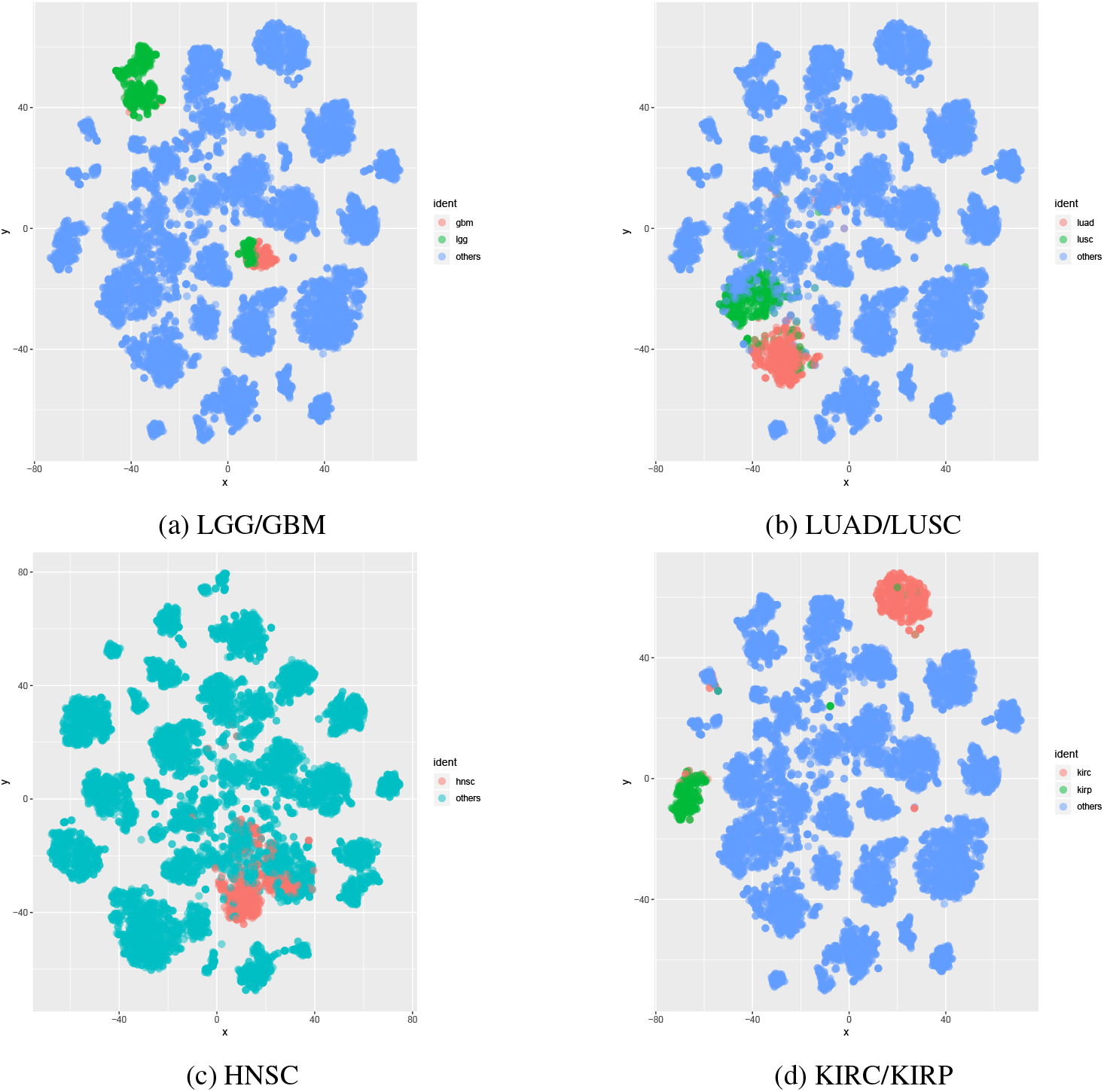
Mapping of 33 types of cancers’ gene expression data with a t-distributed stochastic neighbor embedding (t-SNE), highlighting (S1a) LGG/GBM versus the rest of cancers; (S1b) LUAD/LUSC versus the rest of cancers; (S1c) HNSC versus the rest of cancers; (S1d) KIRC/KIRP versus the rest of cancers. Renal cell carcinomas (i.e. KIRC and KIRP) are farther apart from other types of cancers, and also more heterogeneous within, thus information transfer from other cancers is considered more difficult.

**Table S1:**
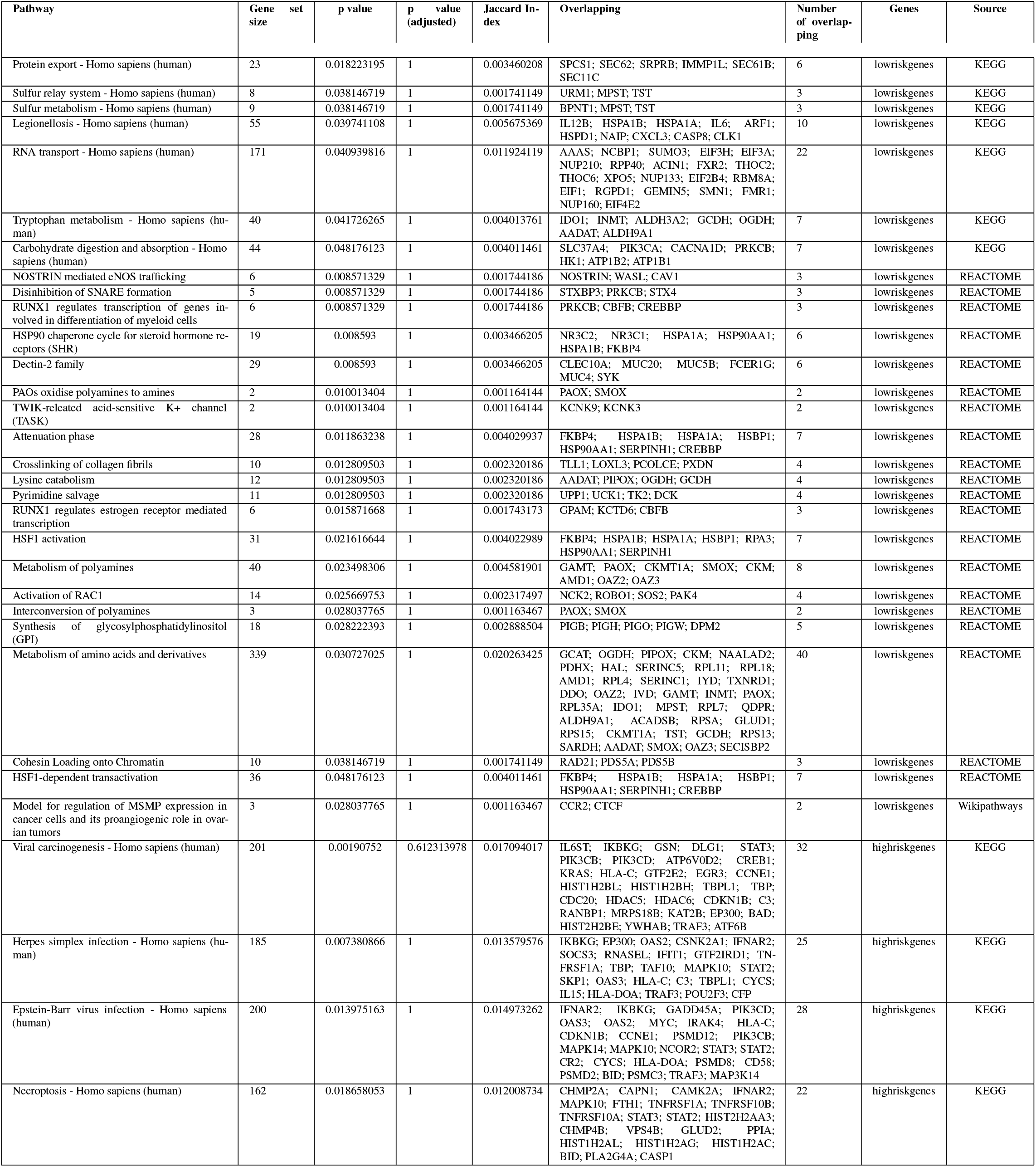
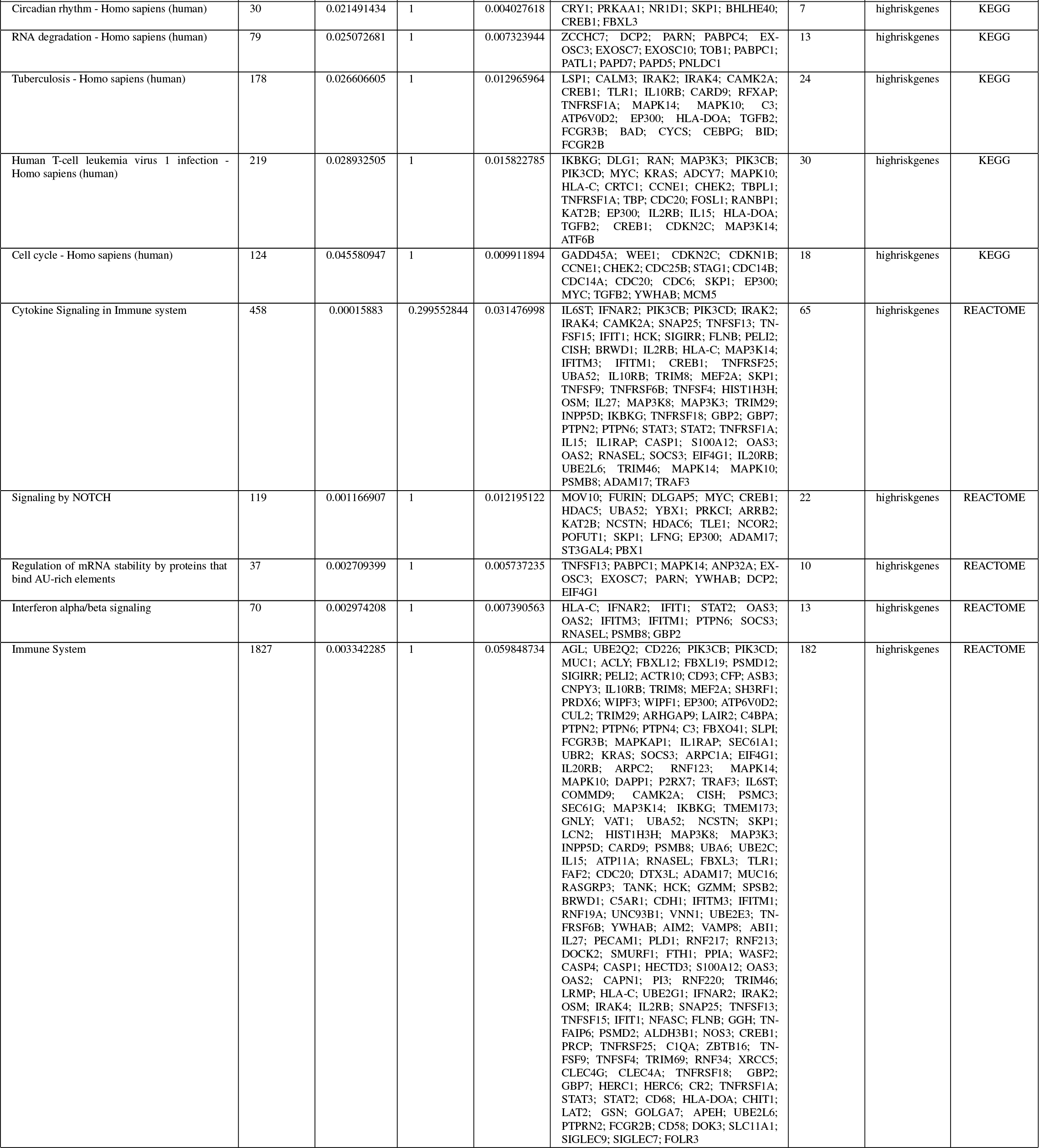
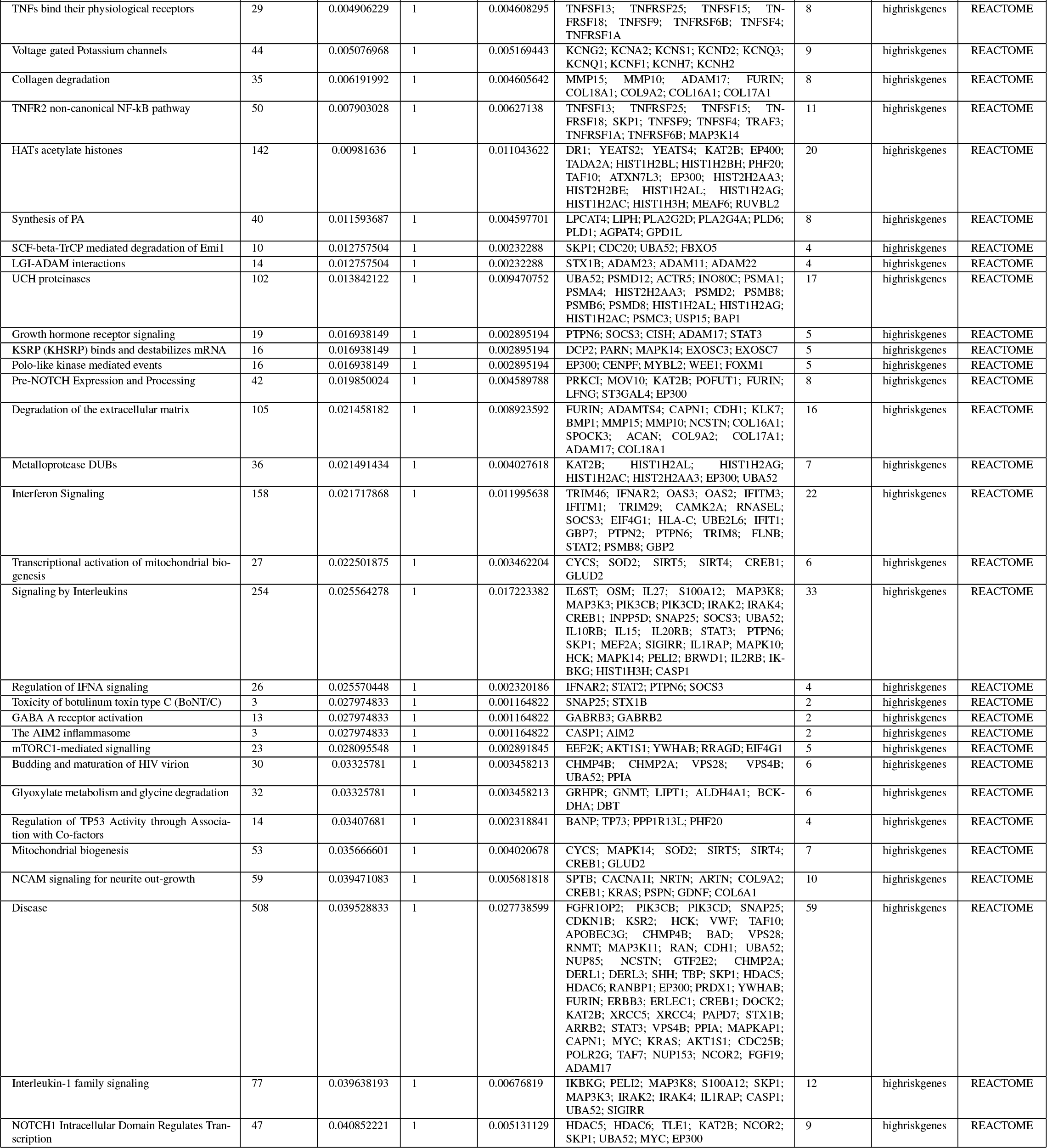
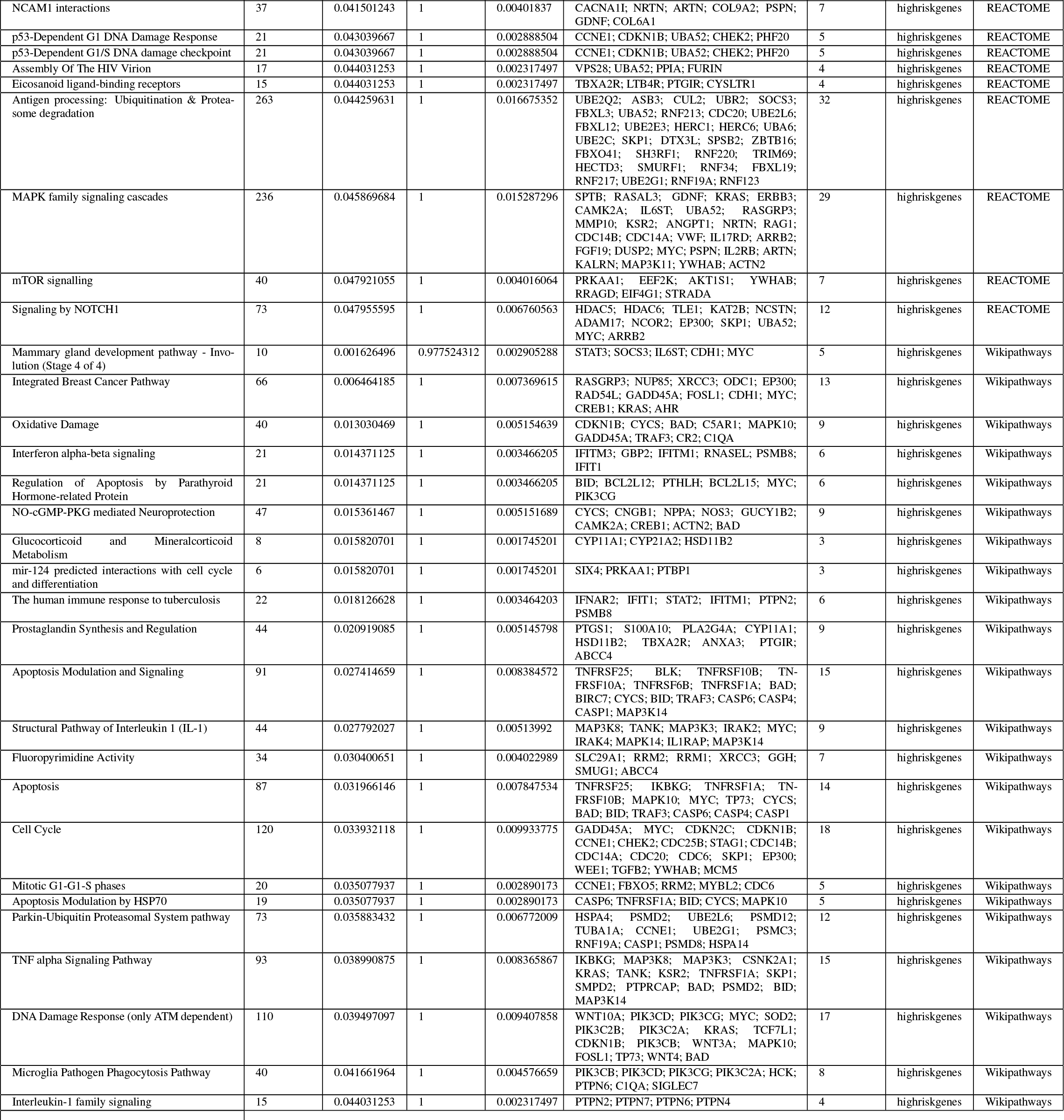
Glioma gene set over-representation

**Table S2:**
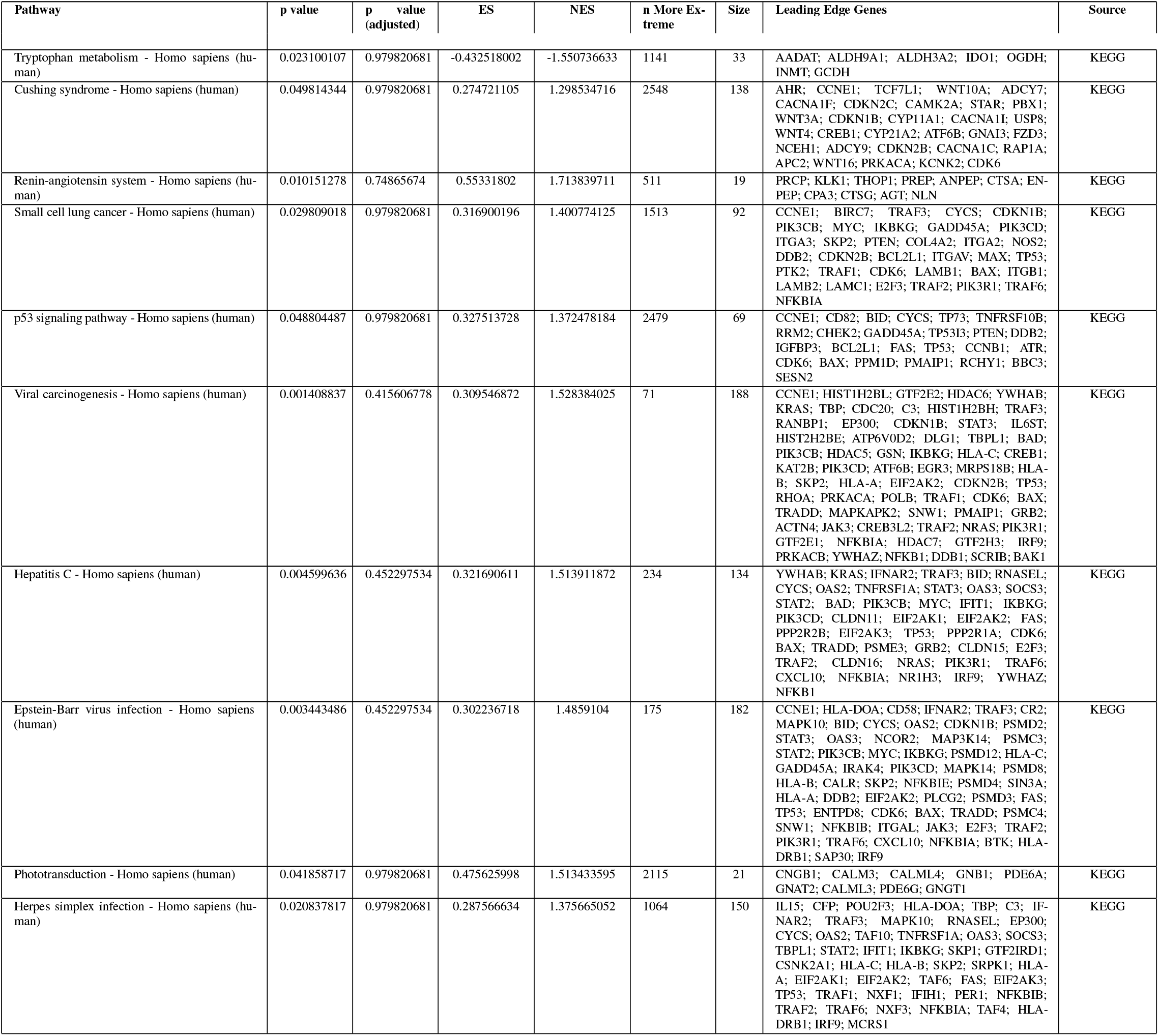
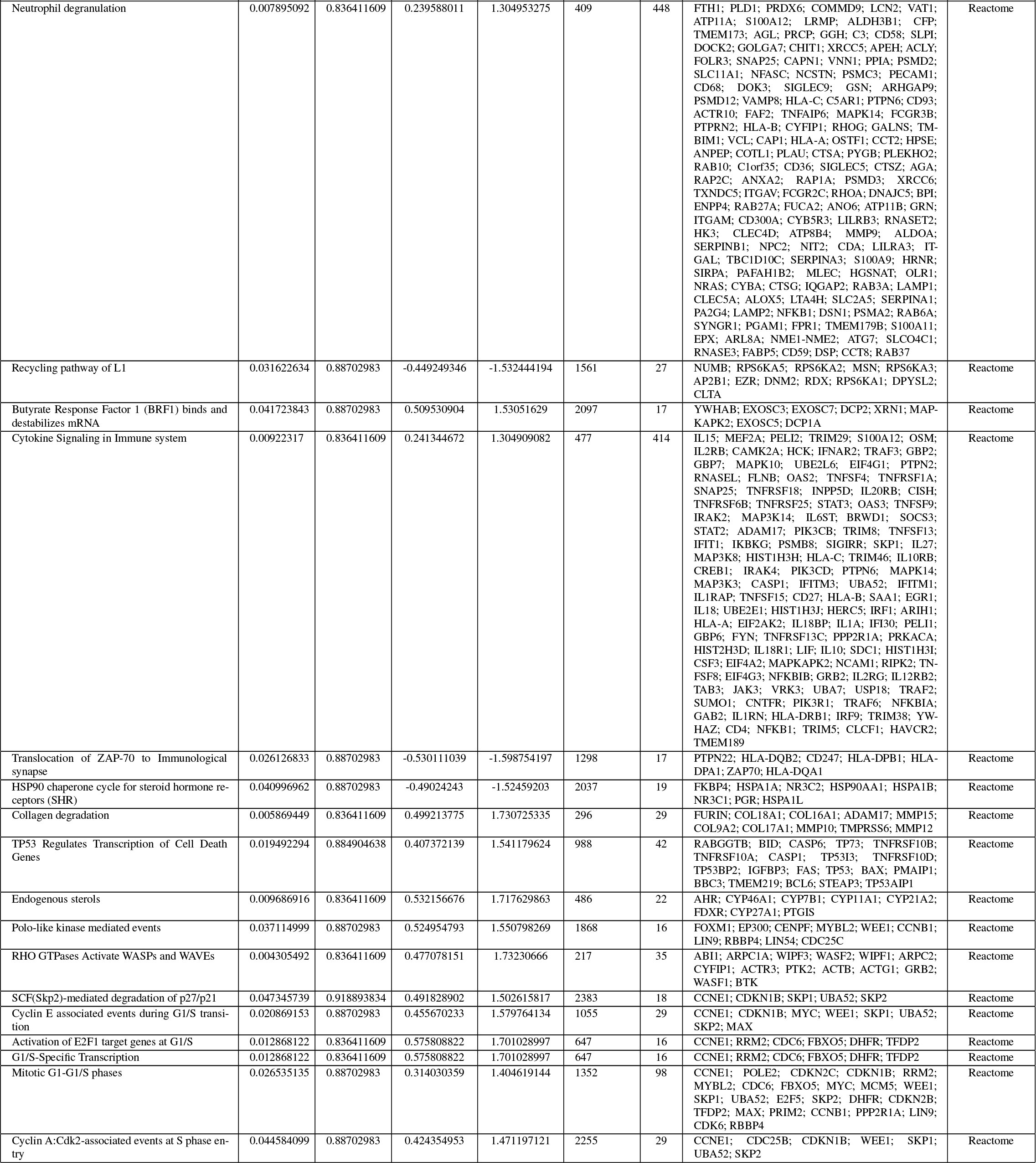
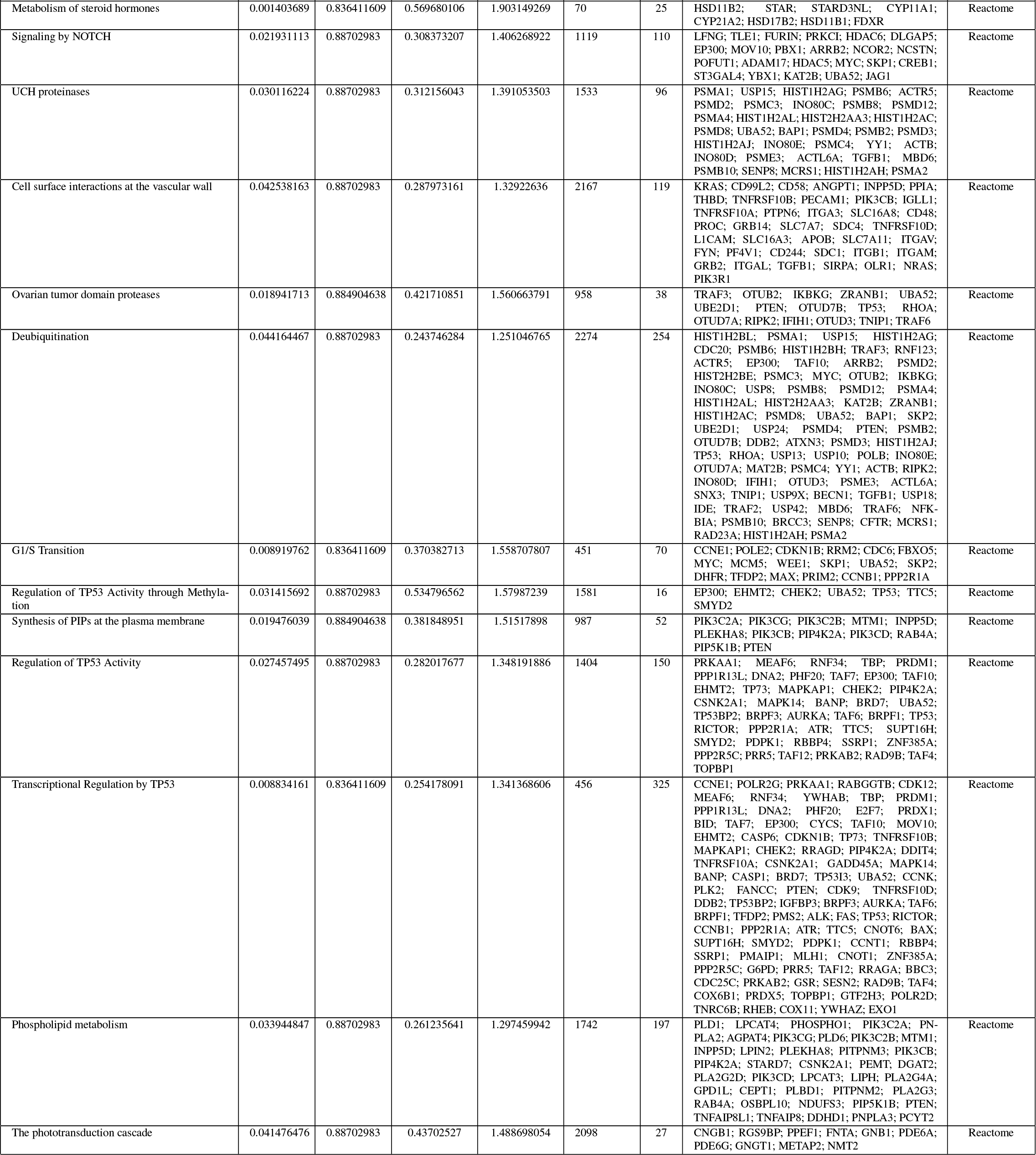
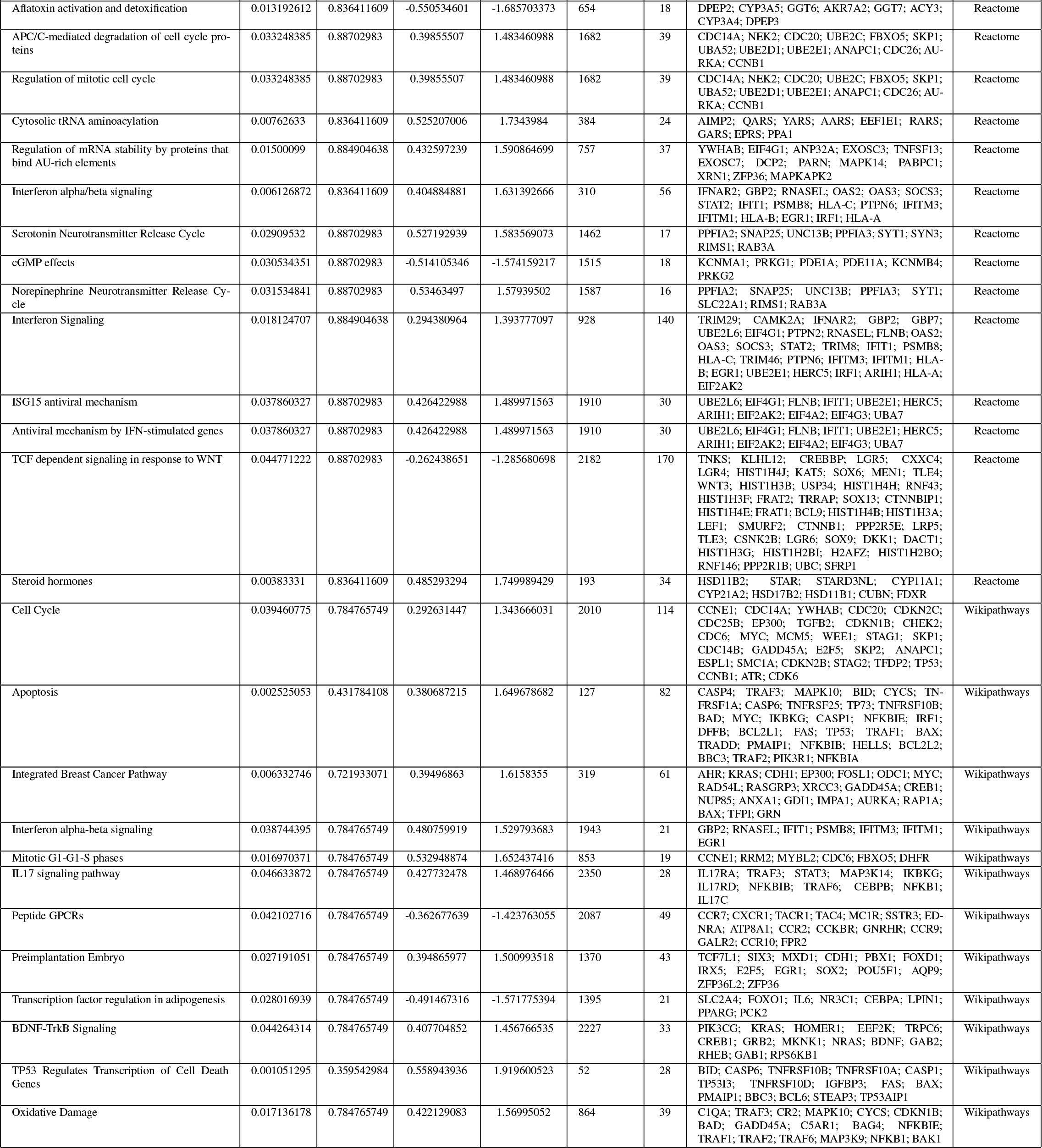
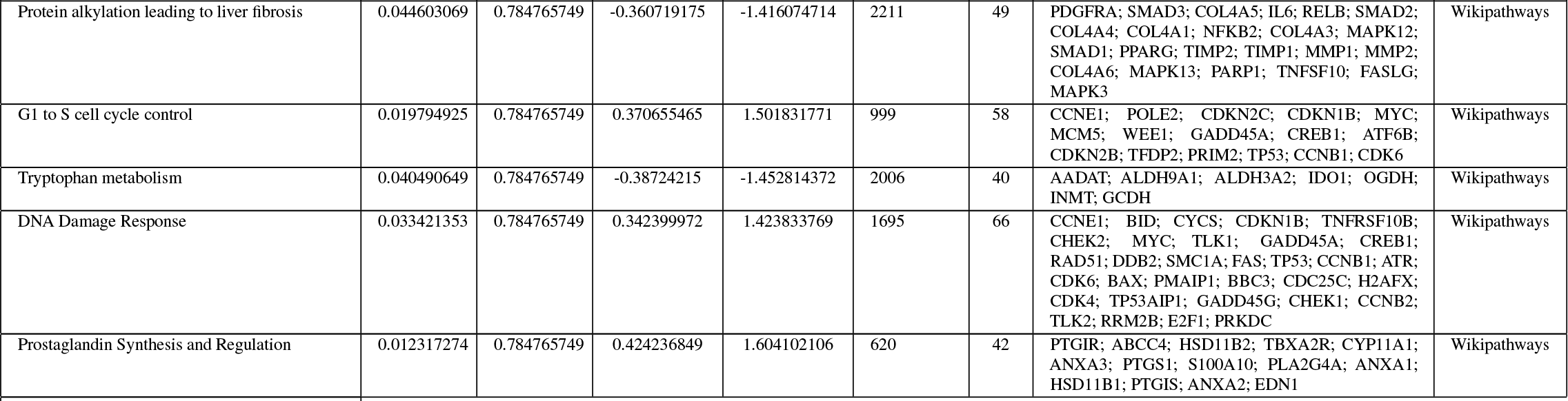
Glioma gene set enrichment analysis (GSEA)

**Table S3:**
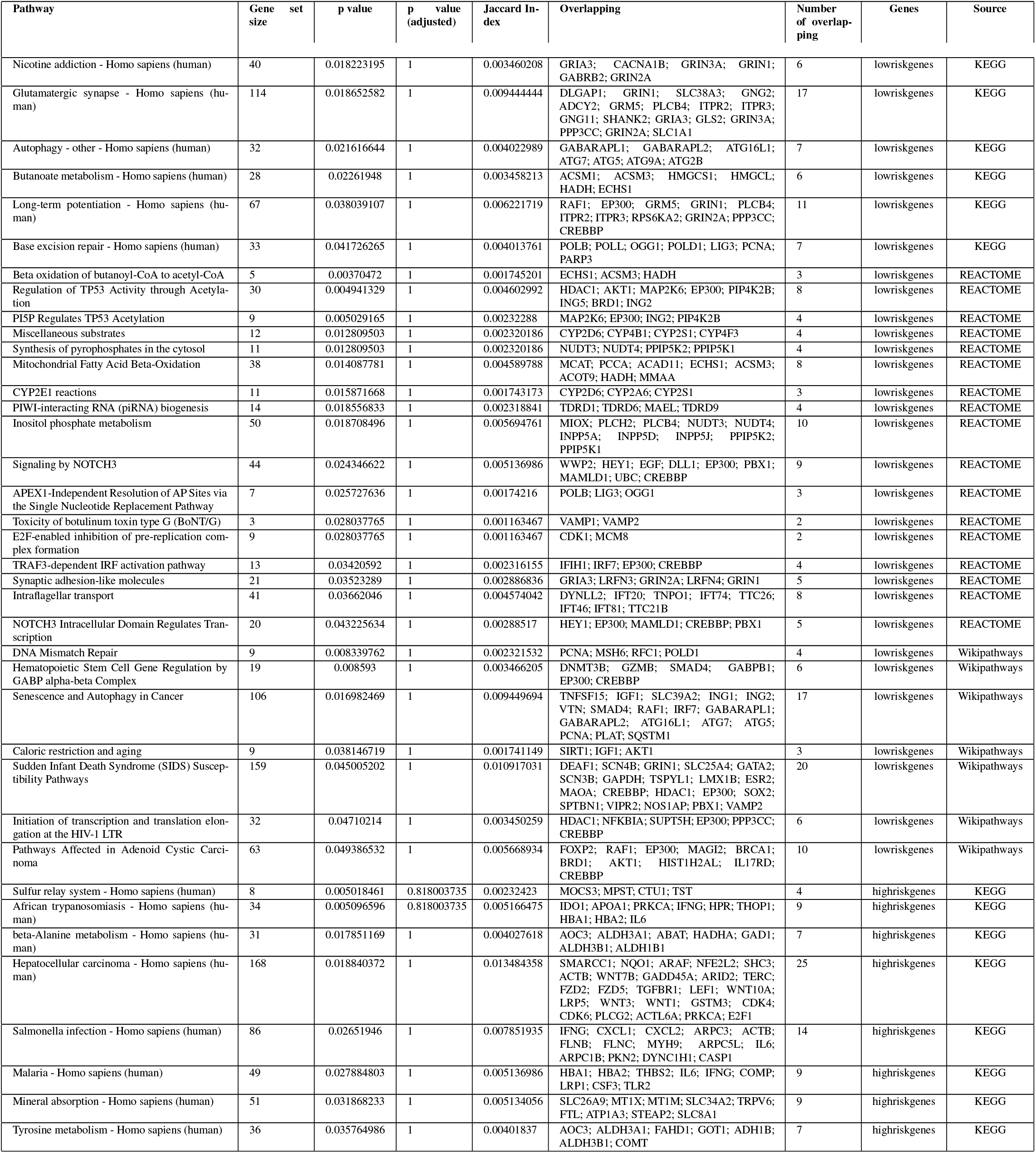
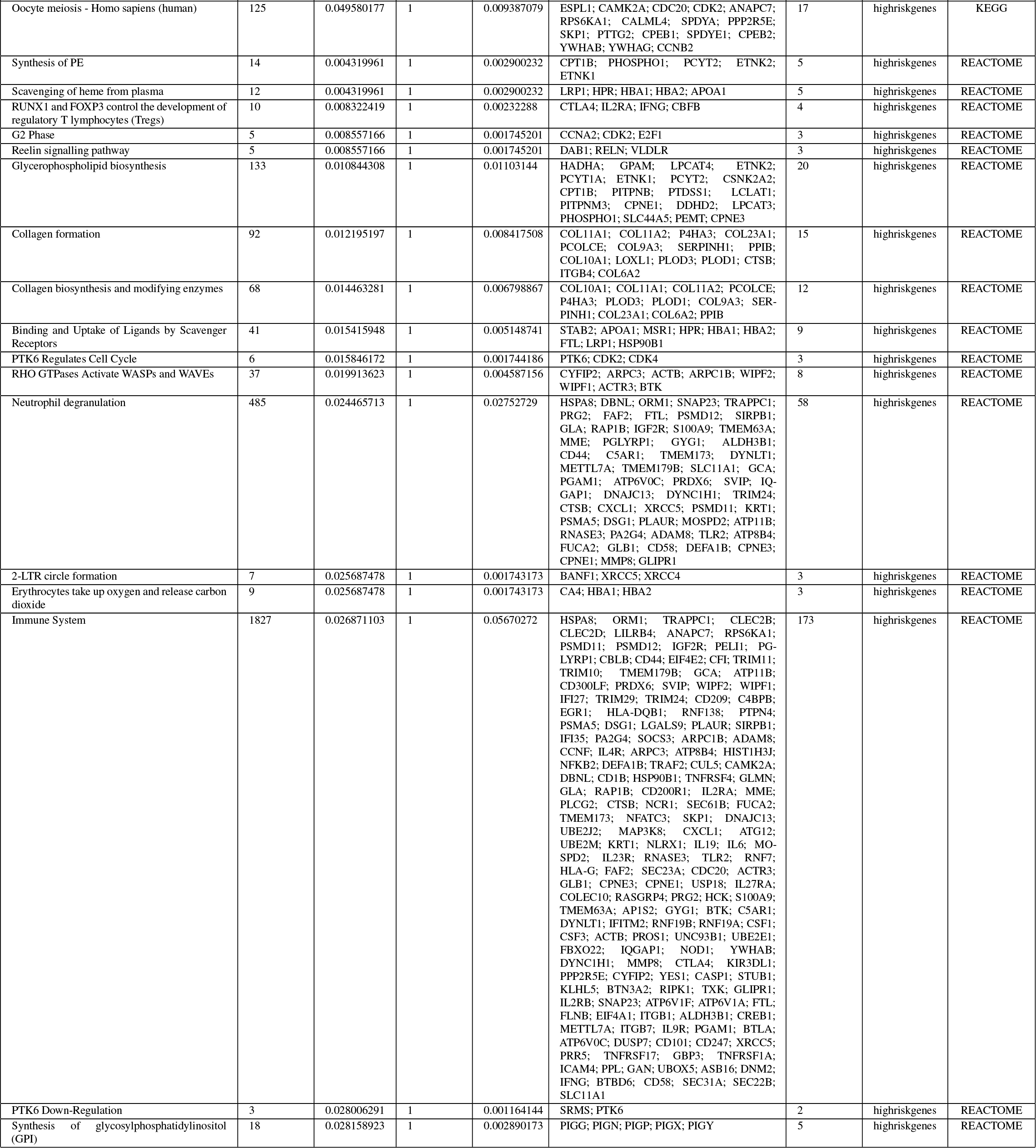
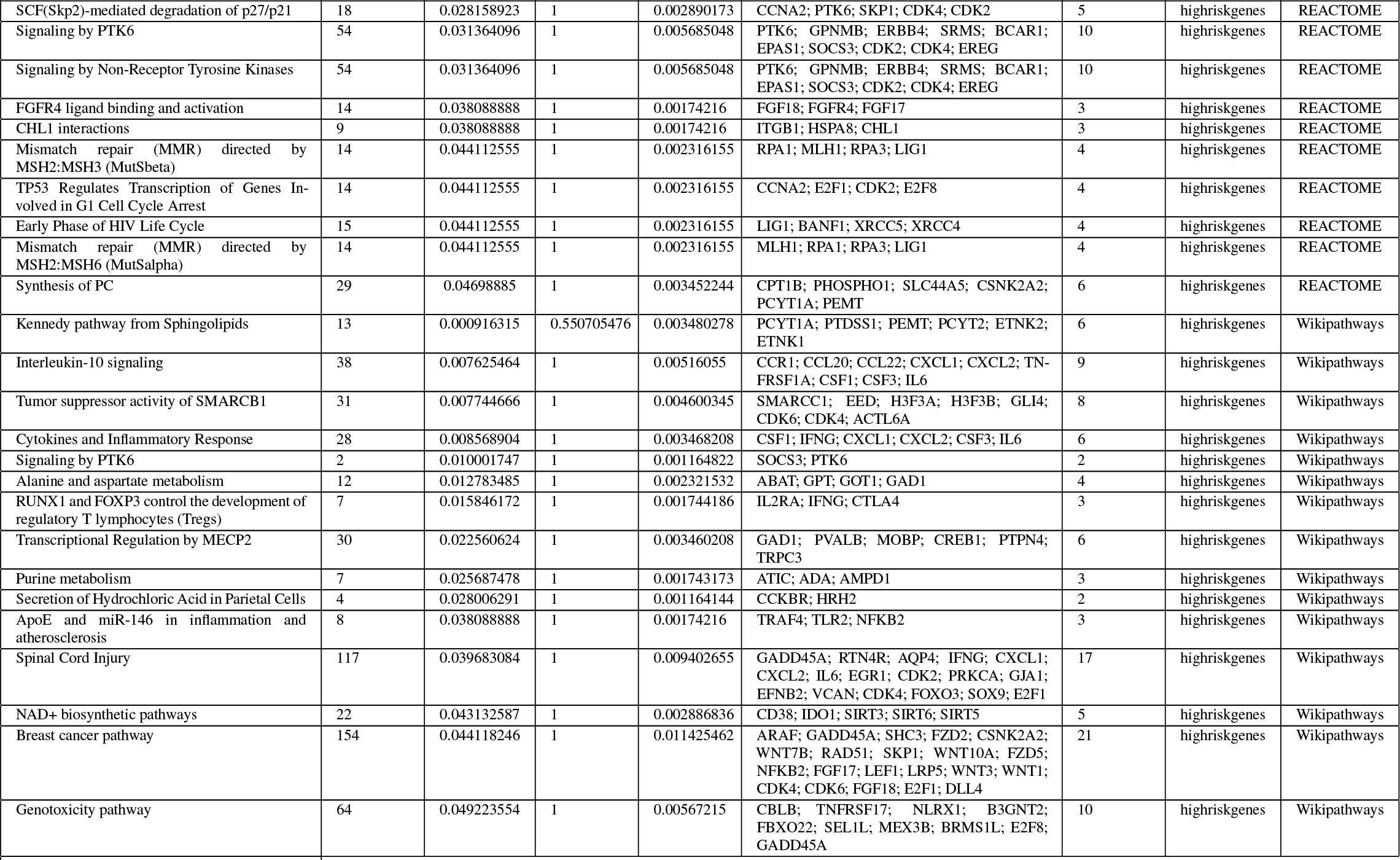
HNSC gene set over-representation

**Table S4:**
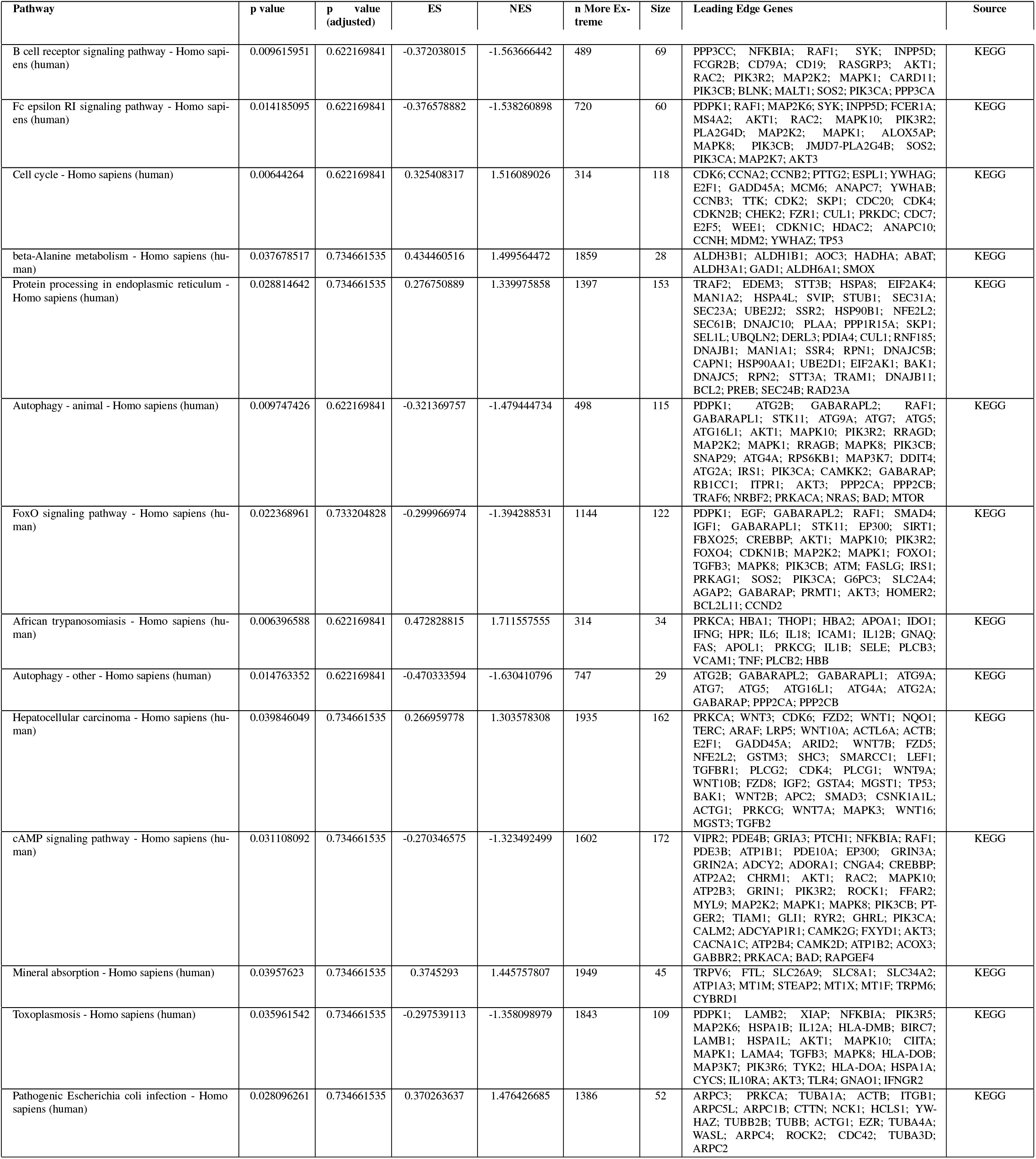
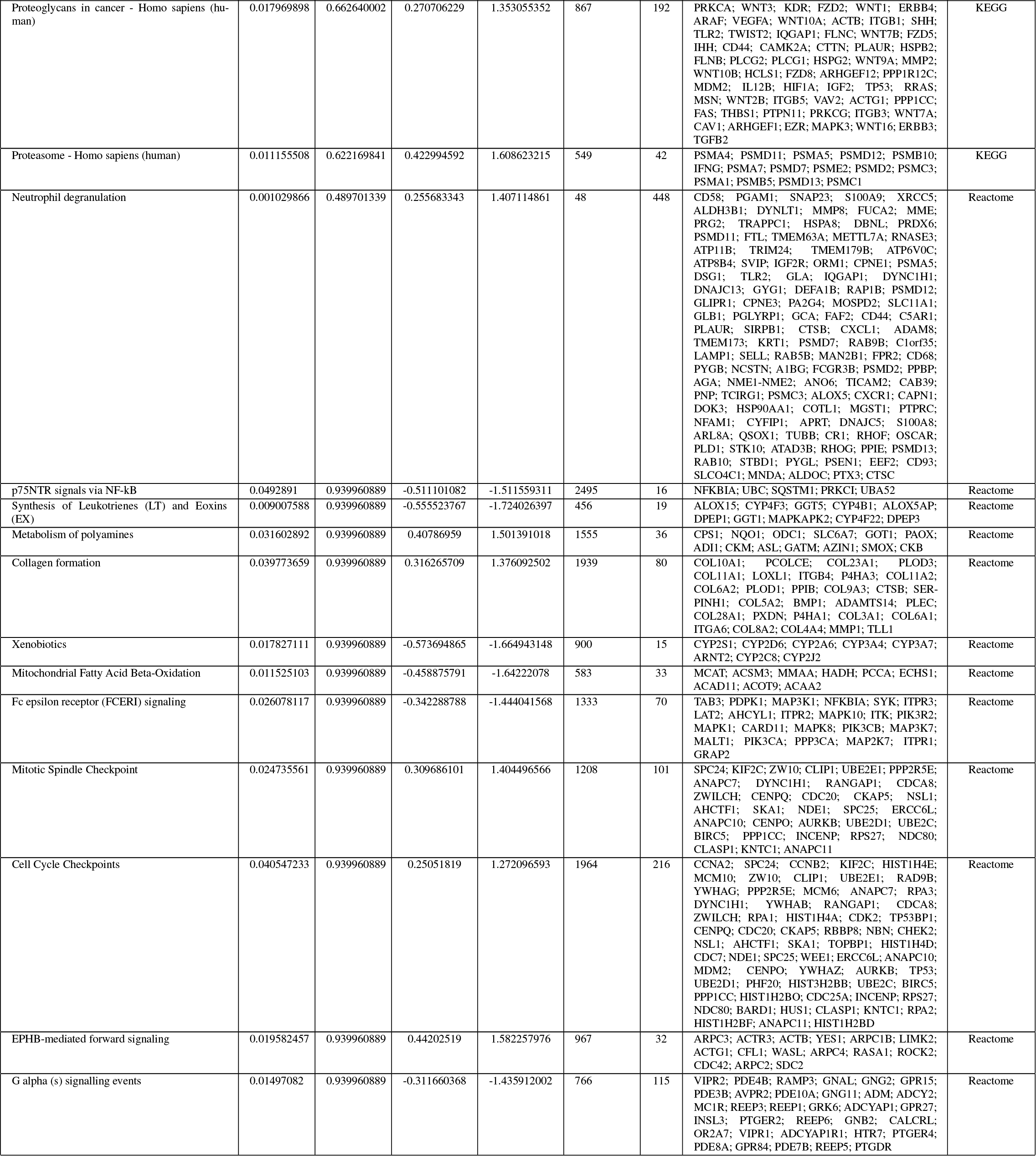
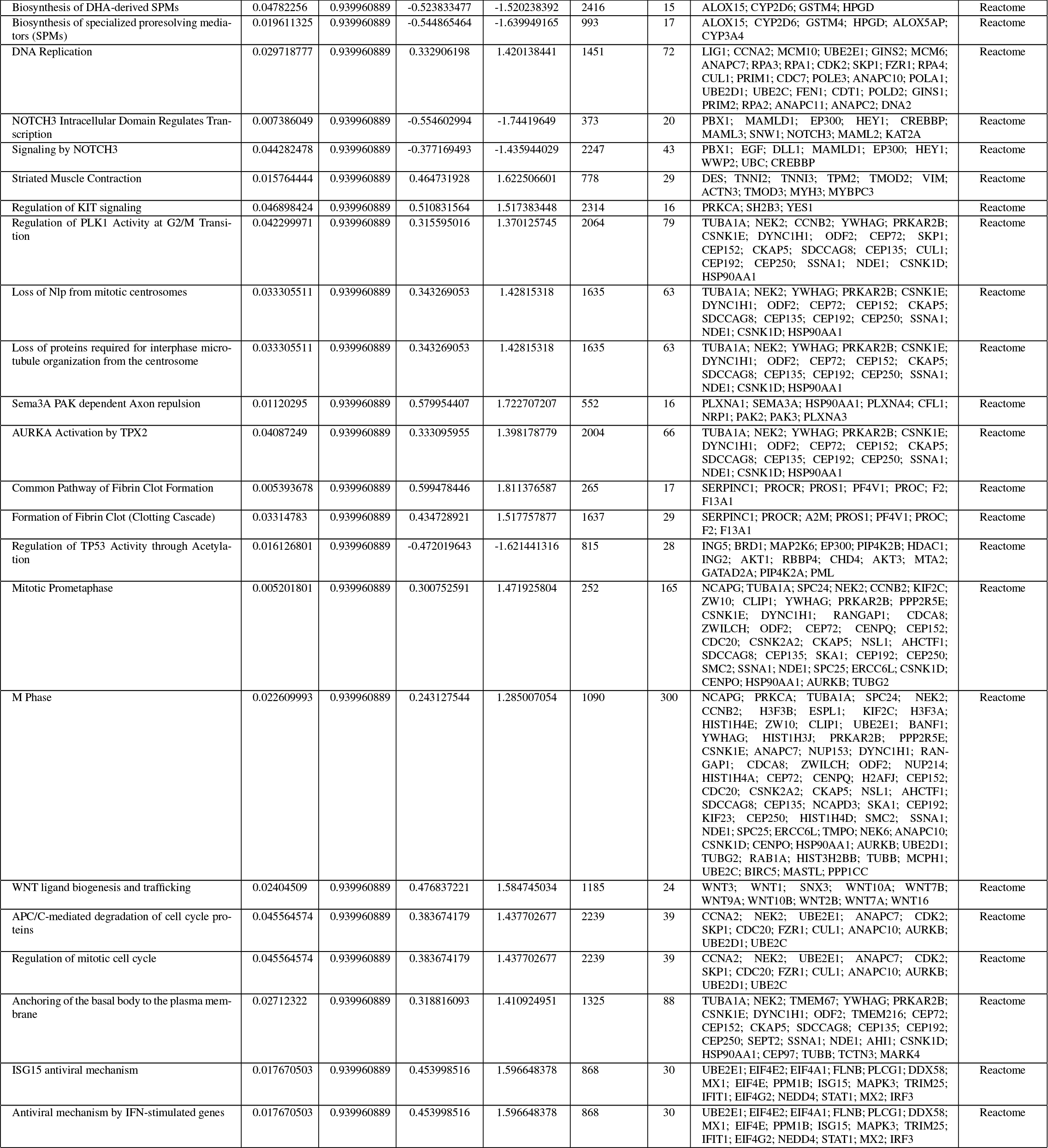
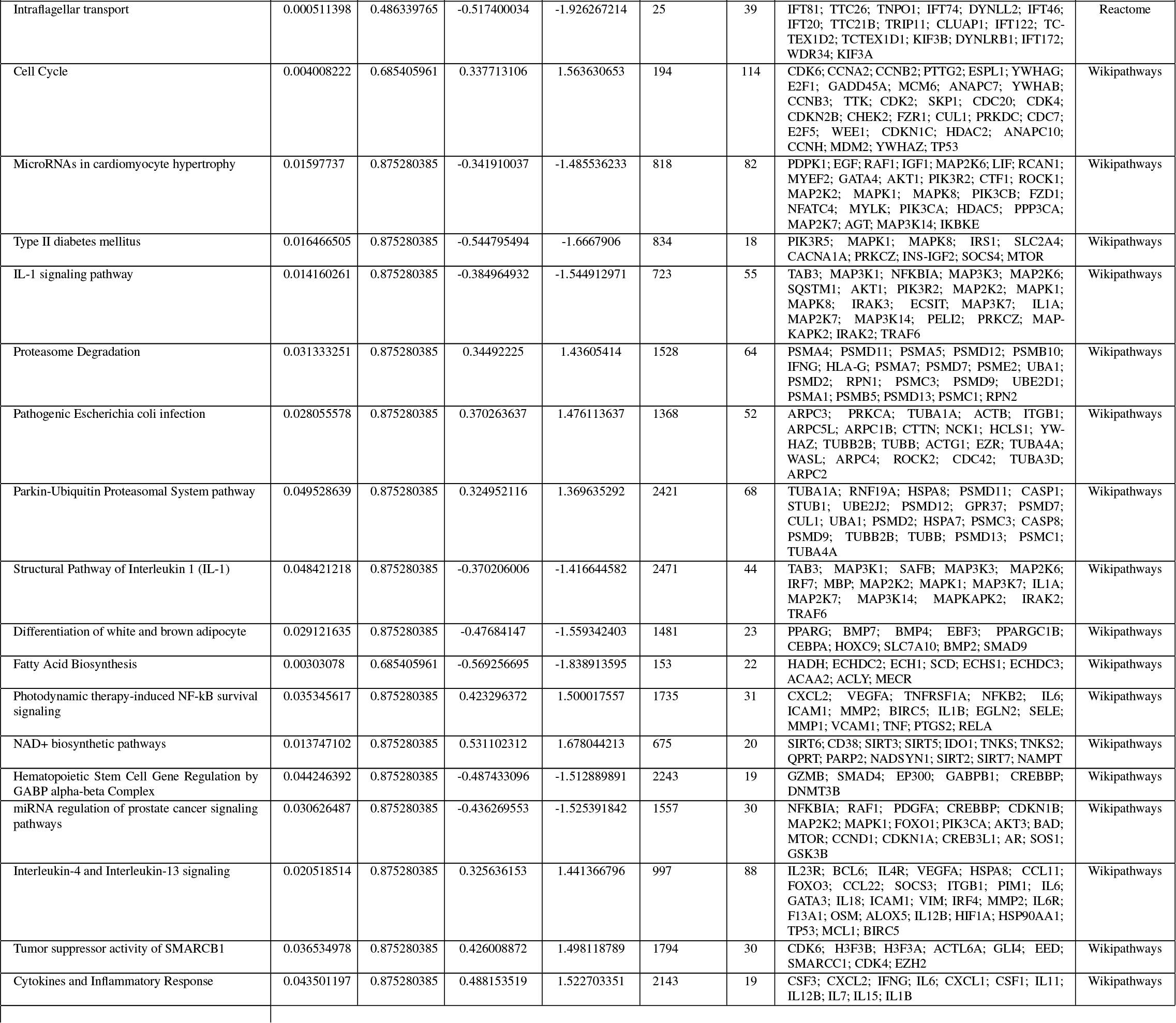
HNSC gene set enrichment analysis (GSEA)

**Table S5:**
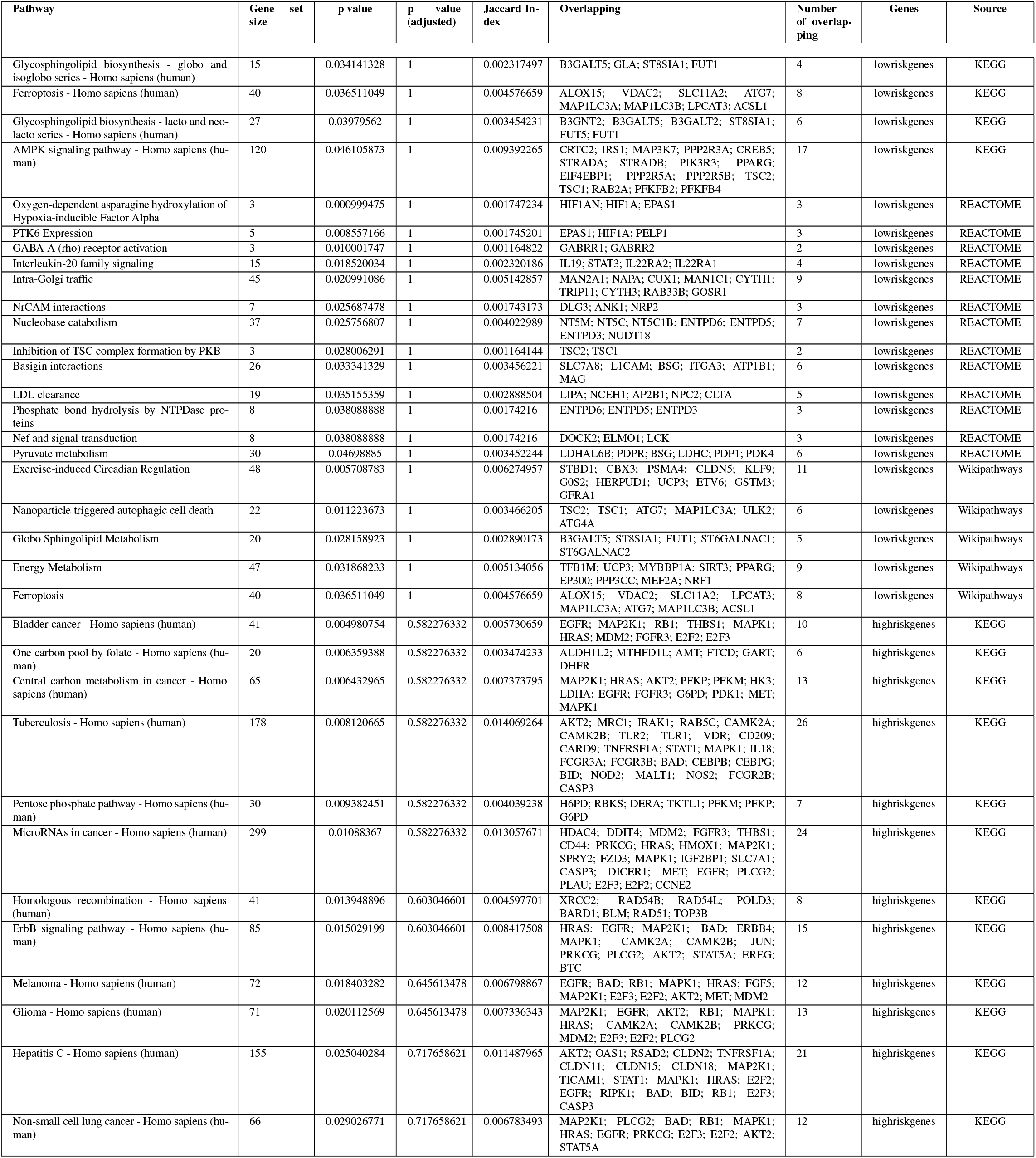
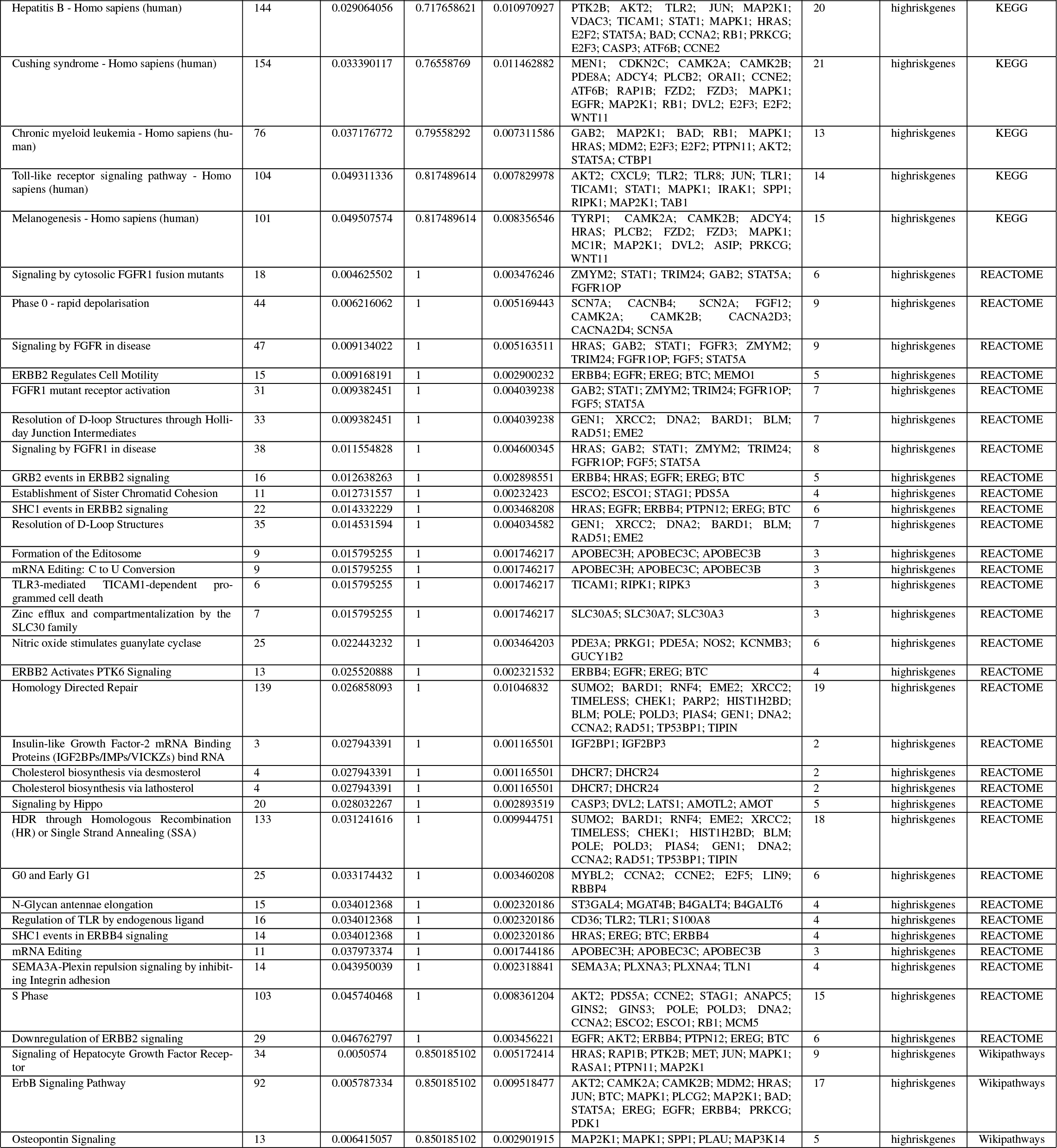
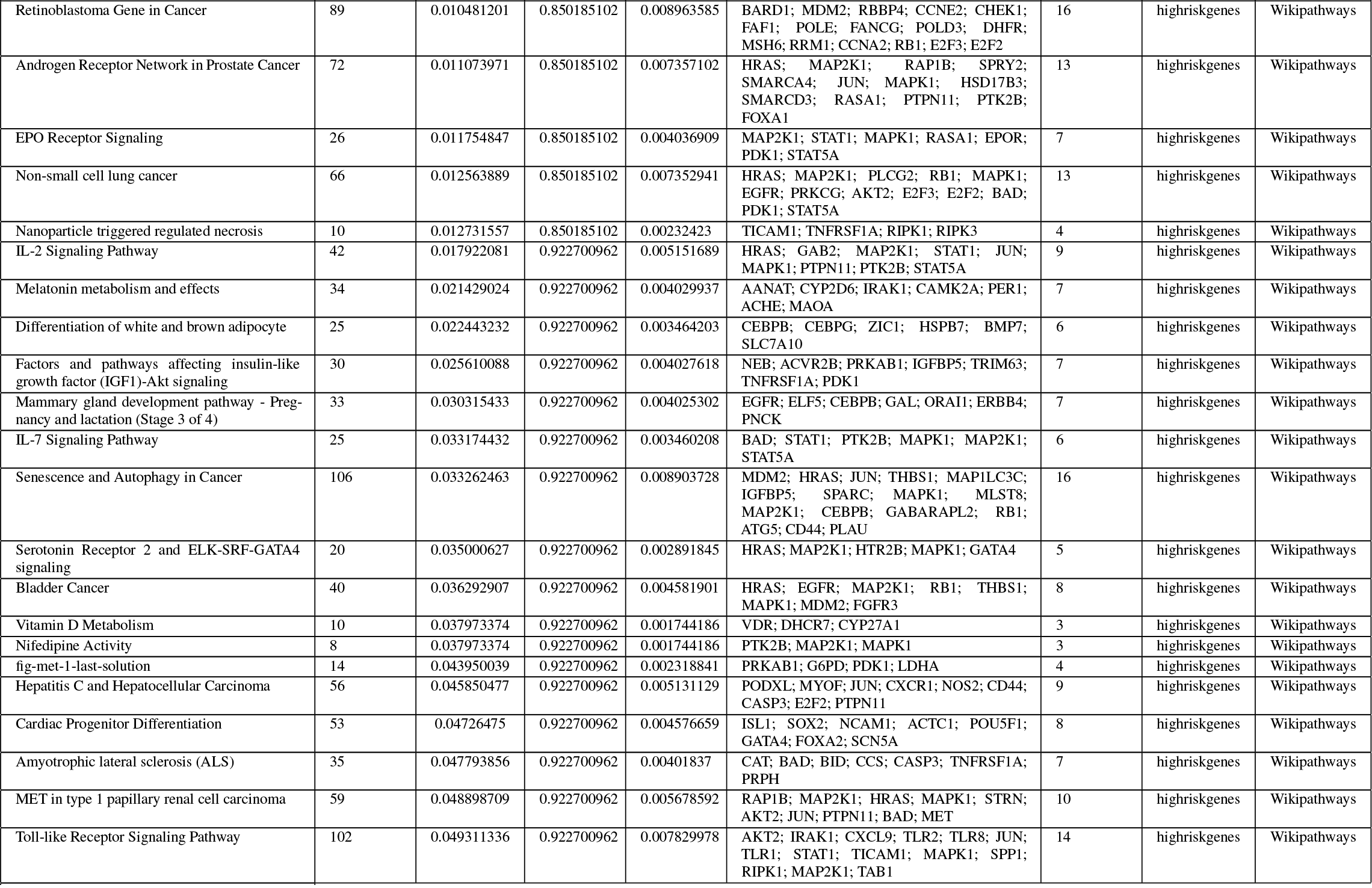
Lung cancer gene set over-representation

**Table S6:**
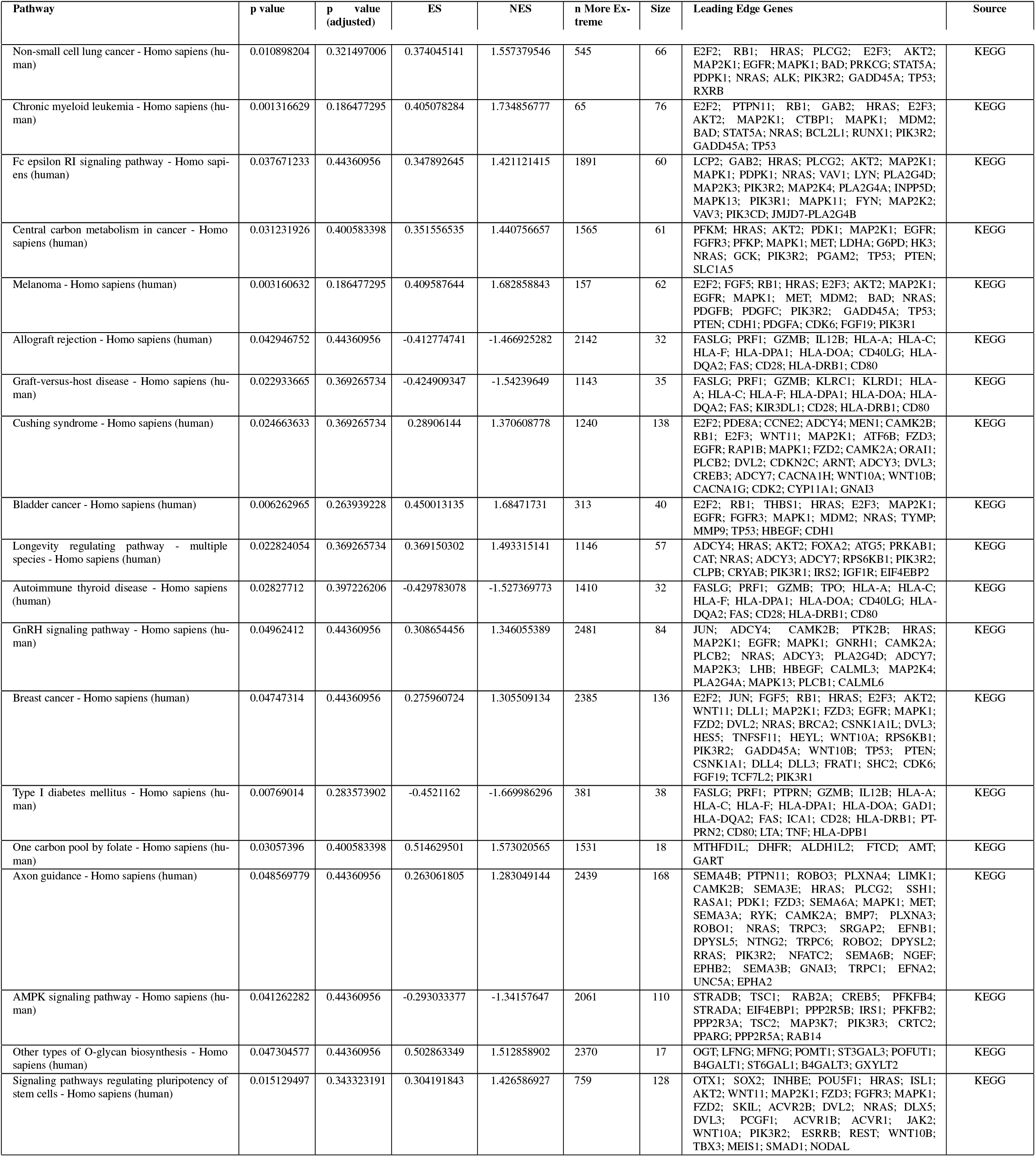
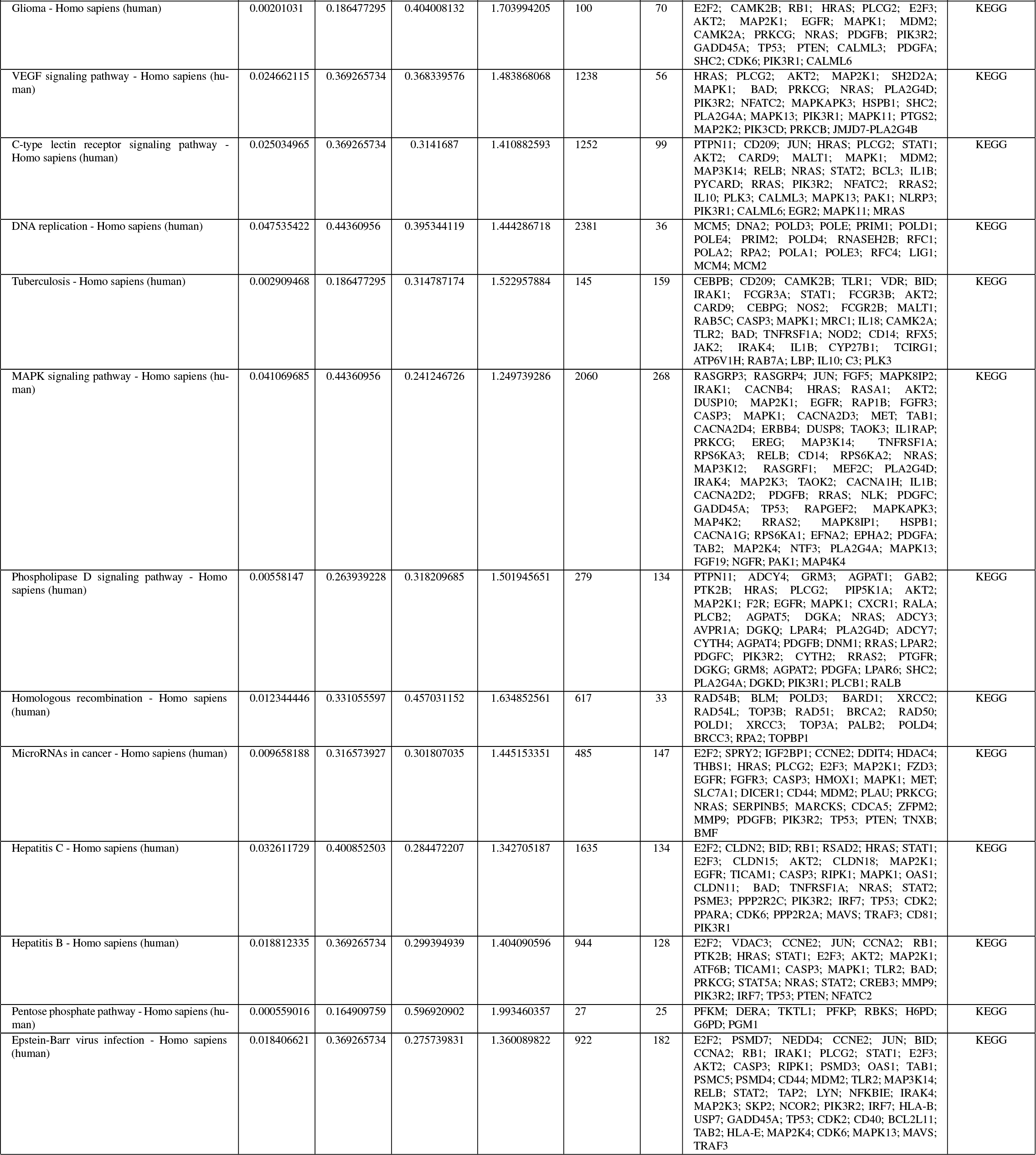
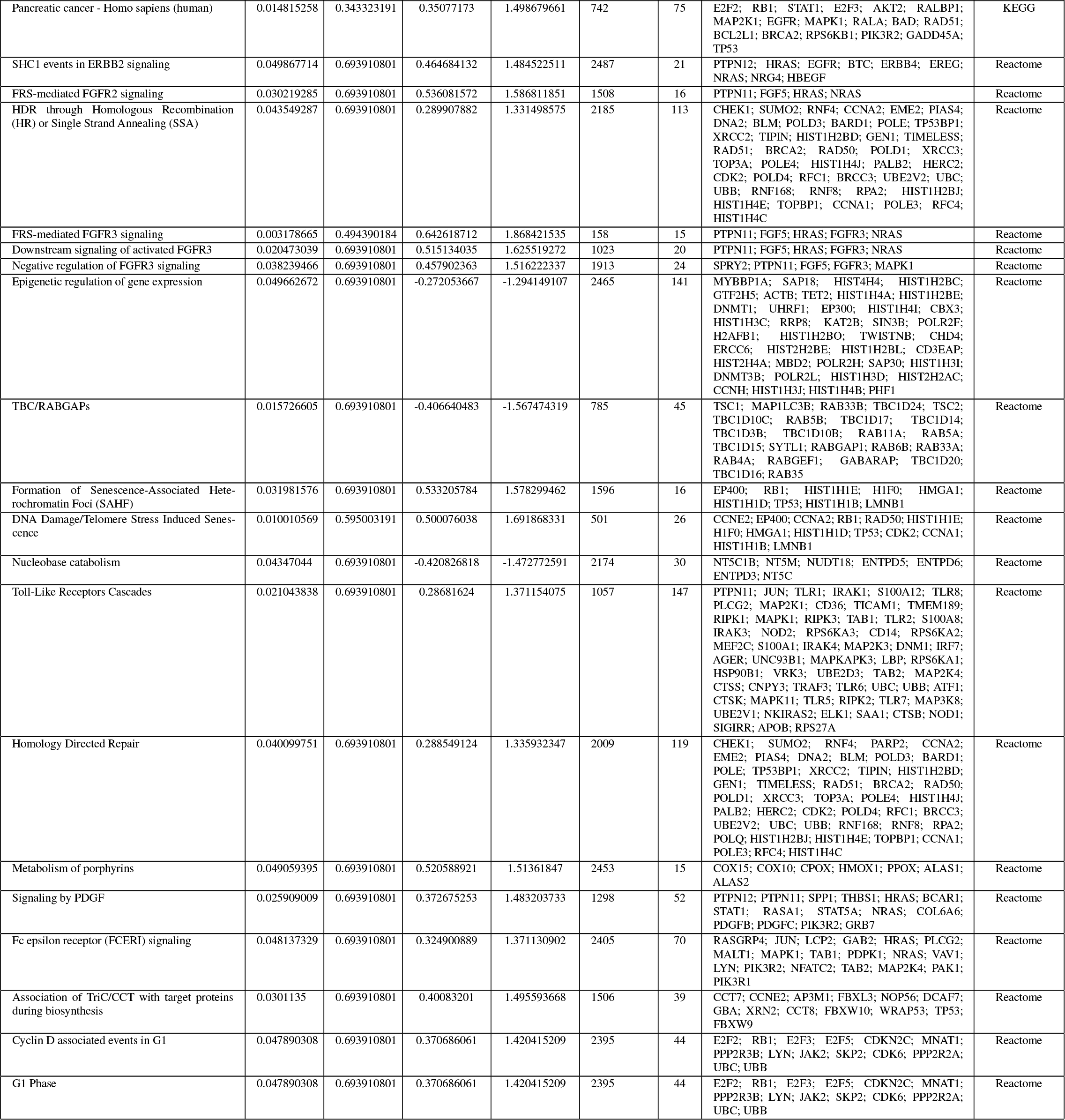
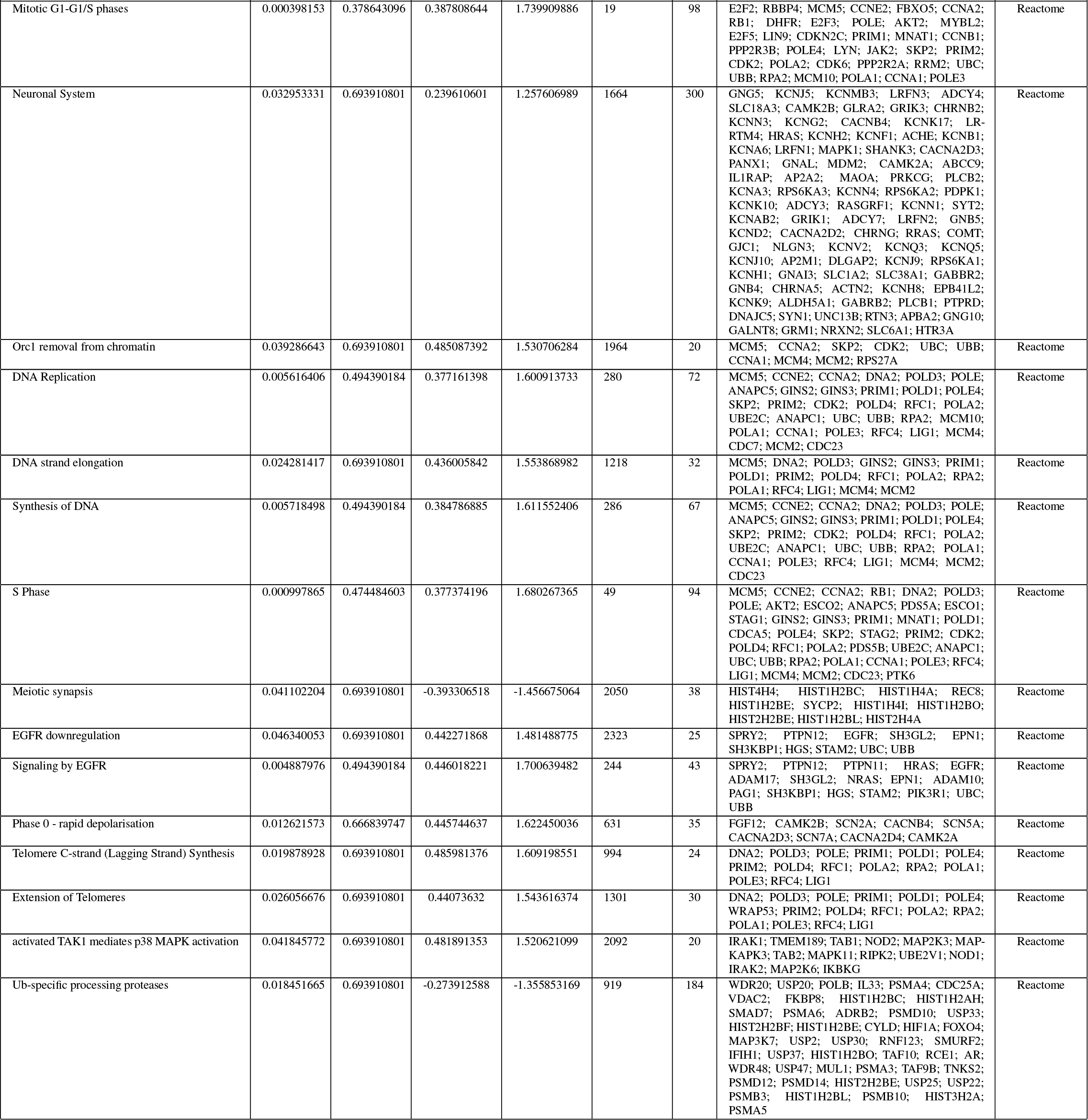
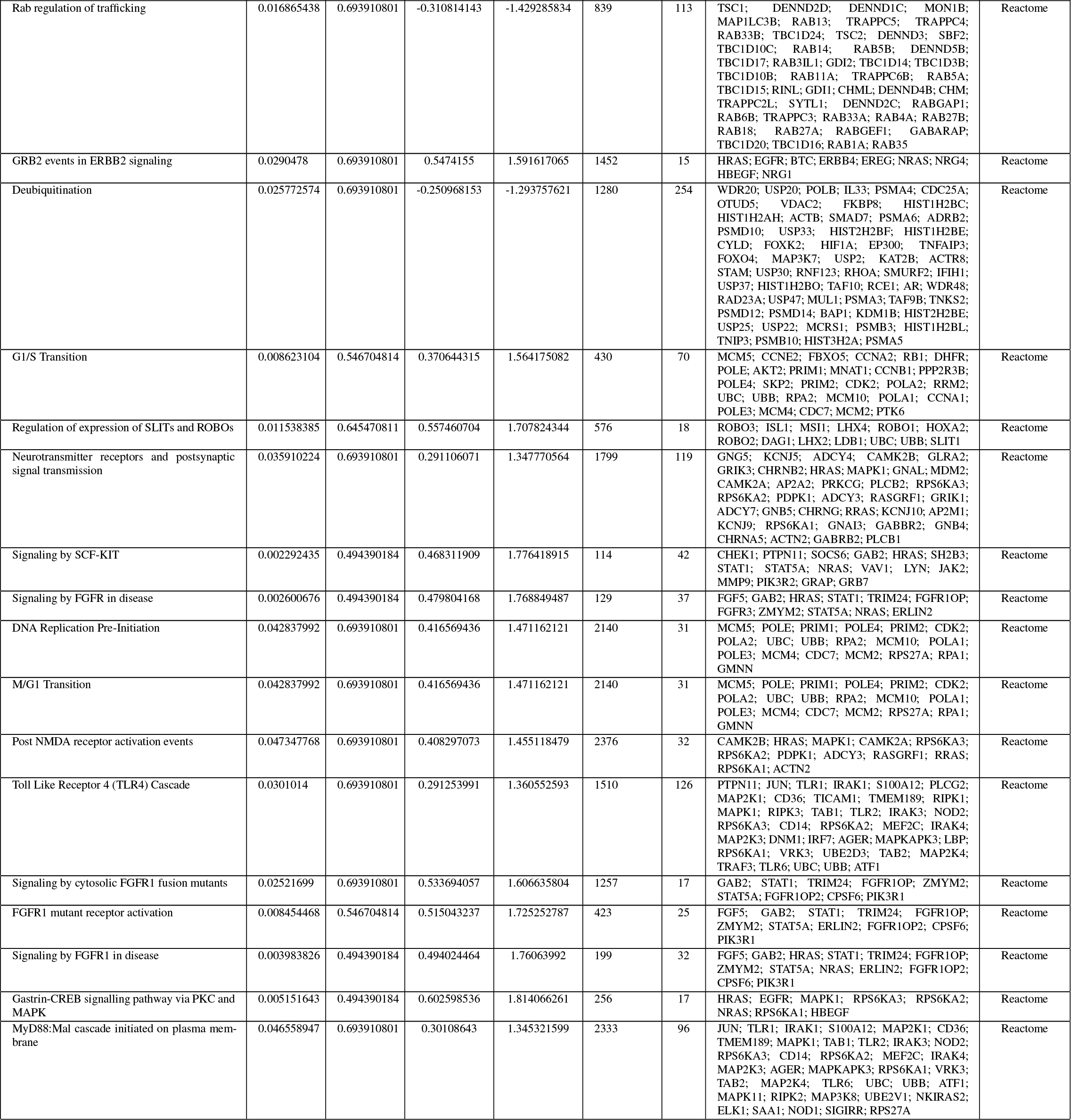
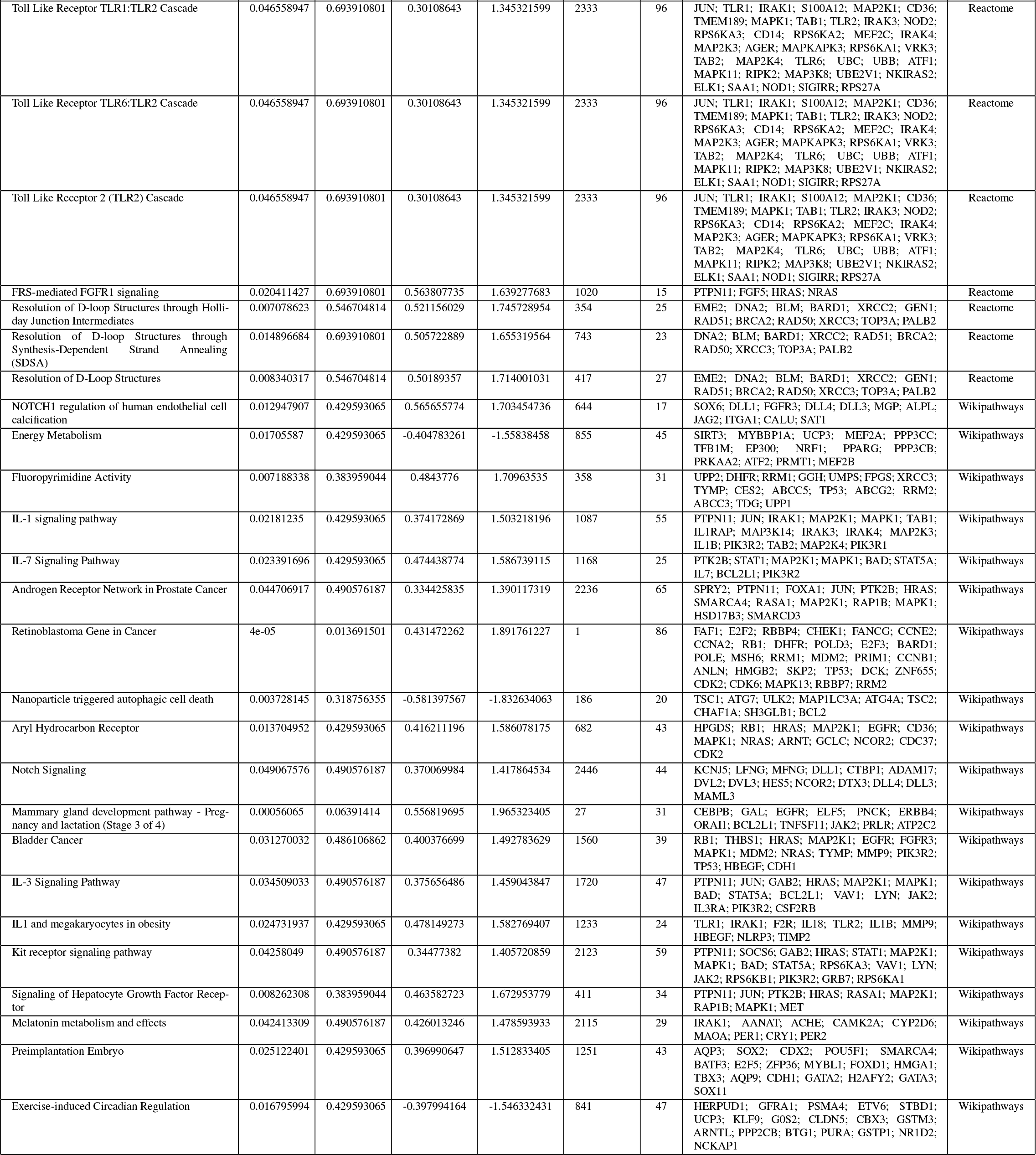
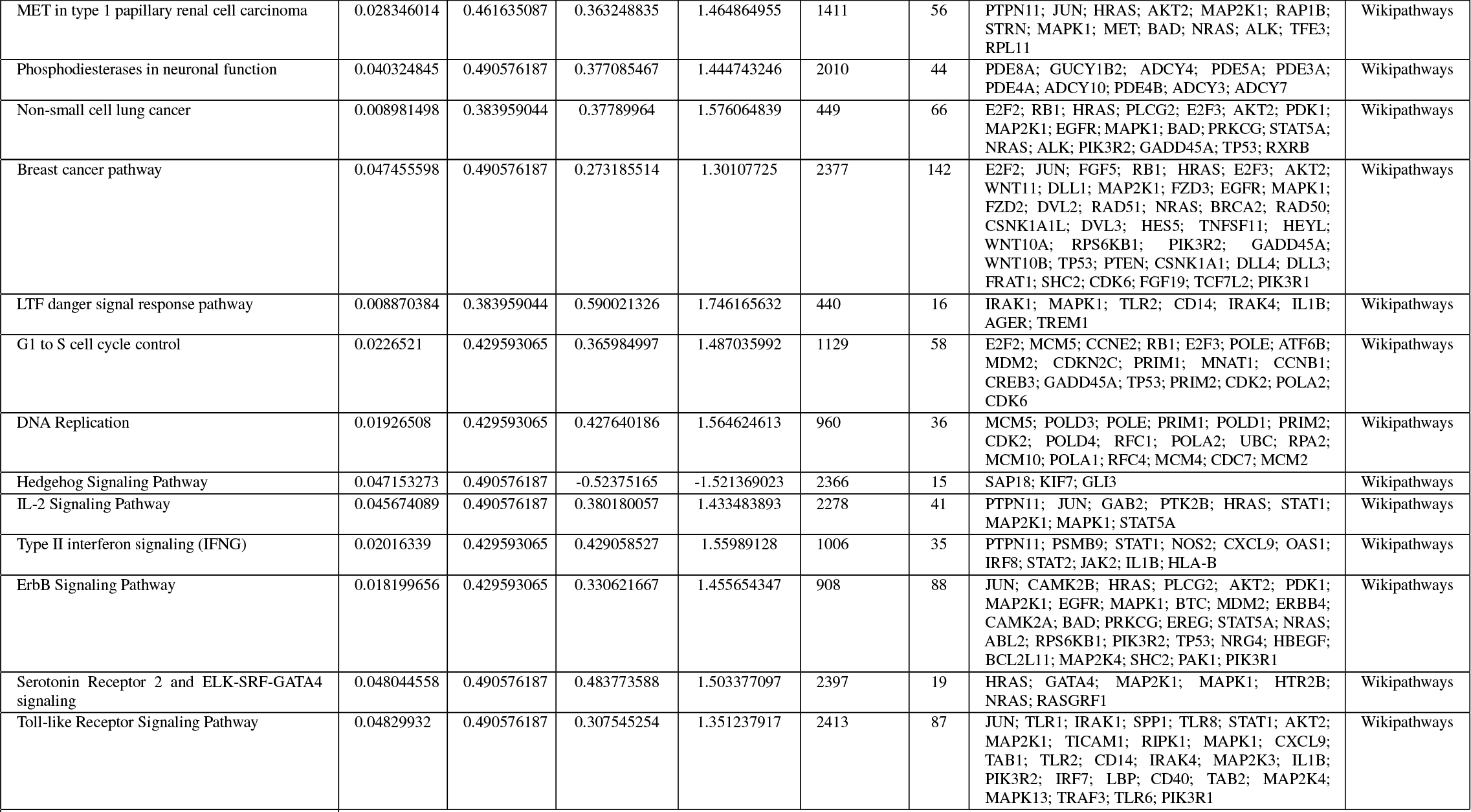
Lung cancer gene set enrichment analysis (GSEA)

